# Spatially resolved single-cell analysis uncovers protein kinase Cδ-expressing microglia with anti-tumor activity in glioblastoma

**DOI:** 10.1101/2023.12.04.570023

**Authors:** Reza Mirzaei, Reid McNeil, Charlotte D’Mello, Britney Wong, Susobhan Sarkar, Frank Visser, Candice Poon, Pinaki Bose, V Wee Yong

## Abstract

Glioblastoma (GBM) is a brain tumor that poses a formidable challenge to treatment options available. The tumor microenvironment (TME) in GBM is highly complex, marked by immunosuppression and cellular heterogeneity. Understanding the cellular interactions and their spatial organization within the TME is crucial for developing effective therapeutic strategies. In this study, we integrated single-cell RNA sequencing and spatial transcriptomics in a GBM mouse model to unravel the spatial landscape of the brain TME. We identified a previously unrecognized microglia subtype expressing protein kinase Cδ (PKCδ) associated with potent anti-tumor functions. The presence of PKCδ-expressing microglia was confirmed in resected human GBM specimens. Elevating tumoral PKCδ expression using niacin or adeno-associated virus in mice enhanced the phagocytosis of GBM cells by microglia in culture and increased the lifespan of mice with intracranial GBM. These findings were corroborated in analyses of the TCGA GBM datasets where low PKCδ samples showed negative pathway enrichment for apoptosis, phagocytosis, and immune signaling pathways, as well as lower levels of immune cell infiltration overall. Our study underscores the importance of integrating spatial context to unravel the TME, resulting in the identification of previously unrecognized subsets of microglia with anti-tumor functions. These findings provide valuable insights for advancing innovative immunotherapeutic strategies in GBM.

## Main

Glioblastoma (GBM) remains the predominant primary malignant brain tumor among adults, posing a significant challenge in the realm of oncology (1). The standard treatment approach involves maximal safe resection using surgery, subsequent radiation, and chemotherapy utilizing the oral alkylating agent temozolomide (1). However, despite these aggressive interventions and other approaches such as tumor-treating fields, patients still face a median overall survival of less than 21 months (2).

Improving prognosis is a formidable hurdle to overcome for GBM patients and the answer may lie within the intricate landscape of the tumor microenvironment (TME) in the brain. The brain TME exhibits two pivotal features: it is highly immunosuppressive and profoundly heterogeneous, both of which significantly contribute to the dismal prognosis associated with GBM (3–6). Unlike other tissues, the brain TME boasts a distinctive composition characterized by a diverse array of functionally specialized organ-resident cells (7). Although single-cell analysis has substantially deepened our understanding of the intricacies within the TME, it grapples with inherent limitations, particularly concerning the spatial architecture of the TME (8–10). This limitation restrains our ability to comprehensively explore the TME architecture and identify subsets of immune cells based on their precise spatial coordinates. Single-cell analysis, while invaluable, offers only indirect insights into cellular interactions due to the loss of spatial organization information (11). Given that the positioning of immune cells within tumors dictates their functions (12–14), unraveling the TME necessitates a focus on their anatomical locations.

Guided by this insight, our hypothesis posits that the GBM microenvironment manifests a dual framework of both functional and spatial organization, with spatial architecture intricately shaping the phenotypes of immune cells. Spatially resolved transcriptomics emerges as a new technology that empowers us to unravel cellular interactions and organization in situ, thereby deciphering the intricate ecosystem of GBM. To validate our hypothesis, we conducted comprehensive analyses, employing both single-cell RNA sequencing (scRNA-seq) and spatial transcriptomics techniques on a GBM mouse model. Additionally, we integrated published scRNA-seq data of human GBM and performed immunofluorescent analyses on human specimens, providing a comprehensive exploration of the brain TME. Our results identify a previously unrecognized and spatially distributed microglia (MG) subset expressing protein kinase Cδ (PKCδ). This cell type subset has potent anti-tumor functions including phagocytic activity, and can be stimulated by a repurposed medication, niacin, to stimulate this MG subset and curb GBM growth.

## Results

### Single-cell and spatially resolved transcriptomic profiling of the GBM tumor microenvironment

To develop a comprehensive understanding of the transcriptomic architecture in the brain TME, we employed the use of a rodent model of GBM (**Fig. 1A**). This model involved the intracranial implantation of mouse brain tumor-initiating cells (BTICs) isolated from NPcis mice, which carry mutations in Nf1 and Trp53, into syngeneic recipients (15). We previously confirmed that this model exhibited the prototypic phenotype of human GBM (16). In vivo bioluminescence imaging confirmed tumor growth in mice (**Supp Fig. 1A**). We obtained 11 brain samples from mice four weeks post-tumor implantation and generated single-cell suspensions from nine mice for scRNA-seq. After sorting of live cells using flow cytometry, we proceeded to process them for scRNA-seq utilizing a droplet-based platform (**Supp Fig. 1B**).

**Figure 1.**
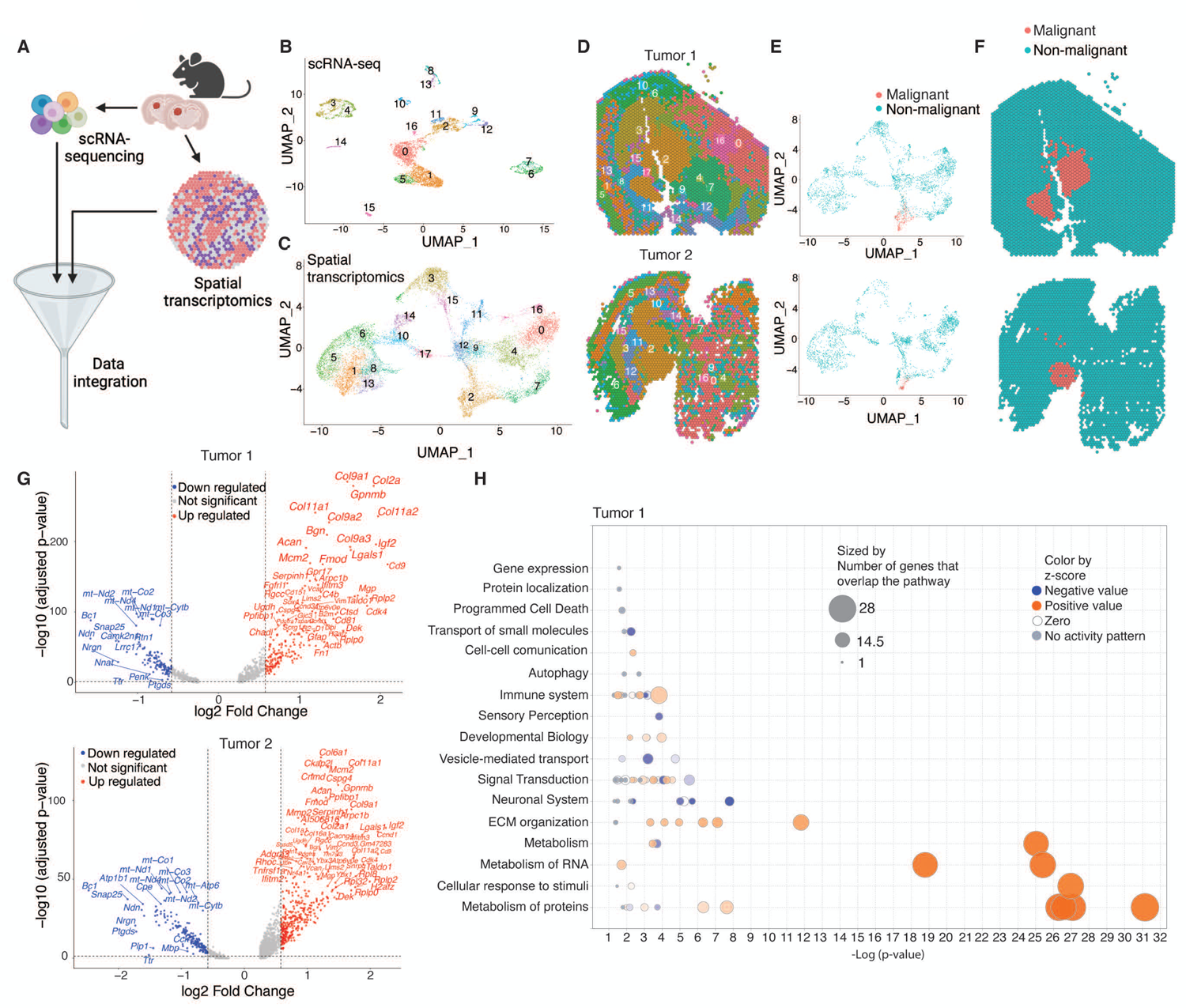
Single-cell and spatially resolved transcriptomic profiling of the GBM tumor microenvironment. A. Schematic representation of the integrated scRNA-seq and spatial transcriptomics analysis conducted in the GBM mouse model. B and C. UMAP plots displaying Seurat clusters identified through scRNA-seq of mouse GBM (B) and spatial transcriptomics analysis (C), respectively. D. Spatial Dimplot of Seurat clusters overlaid on tissue images. E. InferCNV analysis of spatial clusters highlighting malignant and non-malignant spots on tissue images. F. Spatial distribution depicting malignant and non-malignant areas. G. Volcano plots illustrating the differential expression of genes between malignant and non-malignant areas. H. Canonical pathways enriched in DEGs between malignant and non-malignant regions in mouse tumor 1.

The transcriptomic profiles of 5440 cells were generated and grouped into 17 unique Seurat clusters based on gene expression profiles (GEPs) (**Fig. 1B**). Next, cell type deconvolution analysis was performed and identified 11 cell types based on differential expression of known cell type marker genes. This included acute neural stem cells (aNSCs), B cells, dendritic cells, erythrocytes, granulocytes, monocyte derived macrophages (MDMs), microglia (MG), natural killer cells, neurons, oligodendrocytes, and T cells (**Supp Fig. 1C**). Following the initial deconvolution, we noticed that cell type designation would encompass multiple default Seurat clusters, and this prompted us to perform a secondary refined deconvolution analysis. We separated the major cell types of monocytes and MG into subpopulations based on Seurat clustering and trajectory analysis using Monocle3 (**Sup Fig. 1D-H**). By employing this approach, we discerned variations in transcriptomic states among cell types along differentiation trajectories utilizing pseudotime values. This enabled the identification of five subsets of monocytes and four subsets of MG based on GEPs. This deconvolution of the immune contexture within the TME of GBM allowed us to pinpoint distinct subsets of immune cells and their impact on tumor biology. Consequently, we identified 21 cell type clusters ready for downstream analysis (**Sup Fig. 1C** and **H**).

The brain tissue from two additional tumor-bearing mice and one healthy brain without tumor implantation were subjected to Visium spatial transcriptomics as described in our previous study (5). After initial data processing, normalization and batch correction across samples, there was 18 default Seurat clusters identified with 12,885 spots ready for analysis (**Fig. 1C, D, Supp Fig. 2A** and **B**). To computationally identify spots containing tumor cells and their spatial locations we identified cells with copy-number variation (CNV) events using the InferCNV package. Notably, Seurat cluster 2 was enriched for CNV events compared to all other clusters, with common CNV events related to GBM with gains at chromosome 7 (17) (**Fig.1E, Supp Fig. 2C** and **D**). Cluster 2 was localized to known tumoral regions (**Fig. 2F**) aligning with the highly proliferative regions observed in H&E-stained slides reported in our previous study (5).

**Figure 2.**
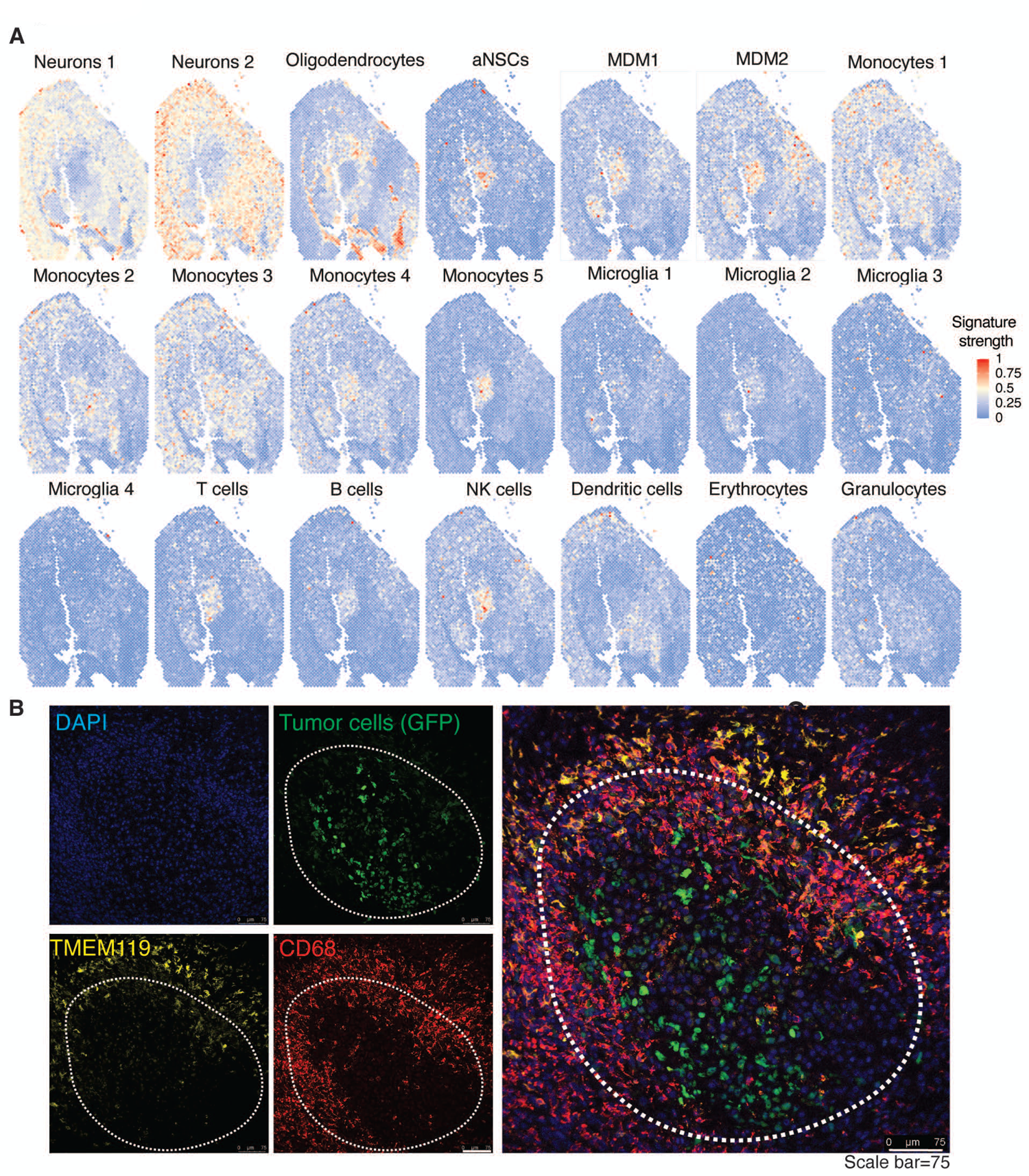
Single-cell spatial heterogeneity in the GBM microenvironment. A. Deconvoluted cell type composition in spatial transcriptomics, determined using cell type-specific expression data obtained from the scRNA-seq dataset of mouse GBM. B. Representative immunofluorescence (IF) image of a mouse brain implanted with syngeneic tumor cells.

Differential expression analysis (DEA) identified differences in GEPs between tumoral spots identified by InferCNV compared to non-tumoral spots within spatial datasets. Volcano plots of differentially expressed genes (DEGs) highlighted statistically significant upregulation of cancer associated genes such as various extracellular matrix (ECM) molecules (*Col6a1*, *Col2a1, Bgn, Acan, Cspg4*, *Vcan, Mgp, Mmp2*), cell cycle genes (*Ccnd3, Cdk4, Ccnd1*), immune-related genes (*Ifitm2, Ifitm3, B2m, Lgals1, H2−D1, Tnfrsf1a, Cd9, Gpnmb*), and complement (*C4b*) in tumoral spots (**Fig.1G** and **Supp Fig. 2E-H**). Ingenuity Pathway Analysis (IPA) of the DEGs from cluster 2 in each mouse sample revealed enrichments in pathways related to metabolism, ECM organization, and the neuronal system, among other significant pathways (**Fig. 1H** and **Supp Fig. 3A**). Additionally, IPA identified notable enrichment of immune system and cell-cell communication pathways in DEGs from cluster 2.

**Figure 3.**
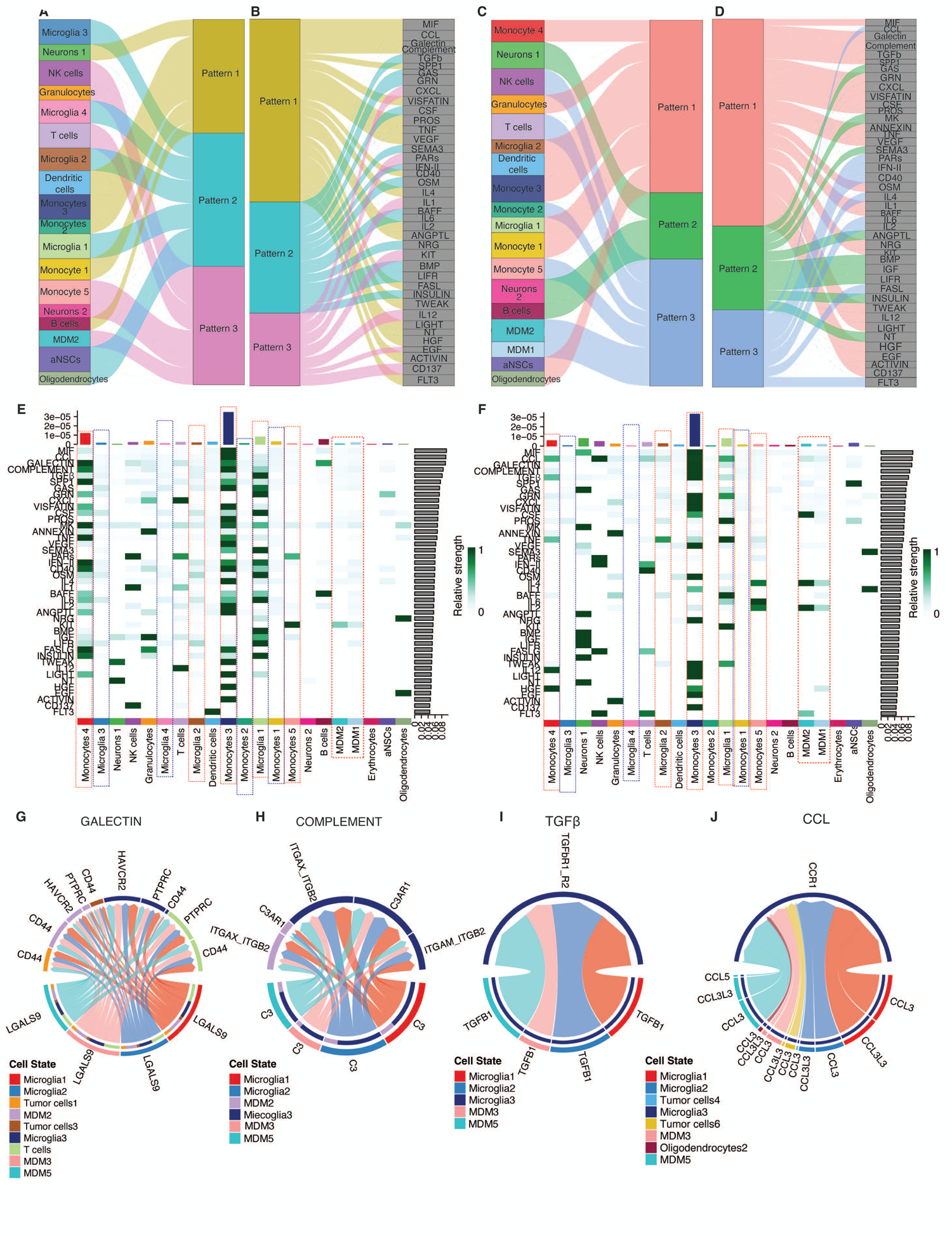
Heterogeneous cellular interactions in the brain tumor microenvironment. A. River plot illustrating patterns of incoming communication in target cells from scRNA-seq data of mouse GBM. B. Signaling networks enriched in three of the incoming communication patterns in mouse GBM. C. River plot displaying patterns of outgoing communication in secreting cells in mouse GBM. D. Signaling networks enriched in three of the outgoing communication patterns. E and F. Heatmaps indicating signals contributing significantly to outgoing or incoming signaling of specific cell groups, respectively in mouse GBM. G-J. Chord diagrams representing all significant signaling pathways from certain cell groups (source cells) to other cell groups (target cells) in human GBMs.

To assess the cellular distribution of the DEGs identified in cluster 2 of the mouse spatial data within the human GBM microenvironment, we analyzed 44,712 cells from a published human scRNA-seq dataset of seven GBM patients (4). After cellular deconvolution as described in our previous study (5) (**Supp Fig. 3B**), we determined the expression profiles of genes from mouse tumoral spots (cluster 2) from spatial data in human scRNA-seq data. We noticed that the ECM genes such as *COL6A1*, *BGN*, *CSPG4*, *VCAN*, and *MGP* are mainly expressed by tumor cells (**Supp. Fig. 3C**). On the contrary, immune related genes such as *IFITM2*, *IFITM3*, *LGALS1*, *TNFRSF1A*, *CD9*, and *GPNMB* exhibited more specific expression by MDMs and MG compared to other cell types in the human GBM microenvironment. This emphasizes the essential role of MG and MDMs in orchestrating the immune response within both the human and mouse GBM microenvironment.

### Single-cell spatial heterogeneity in the GBM microenvironment

To identify the localization and distribution of different cell types with the GBM microenvironment, we performed an integrative analysis using our single cell and spatial transcriptomics datasets. Using the CARD package, we performed a spatially resolved cell type deconvolution using our previously described mouse single cell data as reference. After spatial deconvolution, we identified several patterns of cell type localization related to stromal and immune cell types. High levels of GEP signatures related to neurons were distributed throughout the spatial tissue slides, excluding known tumoral regions (**Fig. 2A** and **Supp Fig. 4A**). High levels of GEP signatures related to oligodendrocytes were also identified, excluding the tumoral regions. Furthermore, aNSC GEP signatures were identified within tumoral regions at the tumor center; this may indicate their pivotal role in the tumor microenvironment related to tumor cell origin.

**Figure 4.**
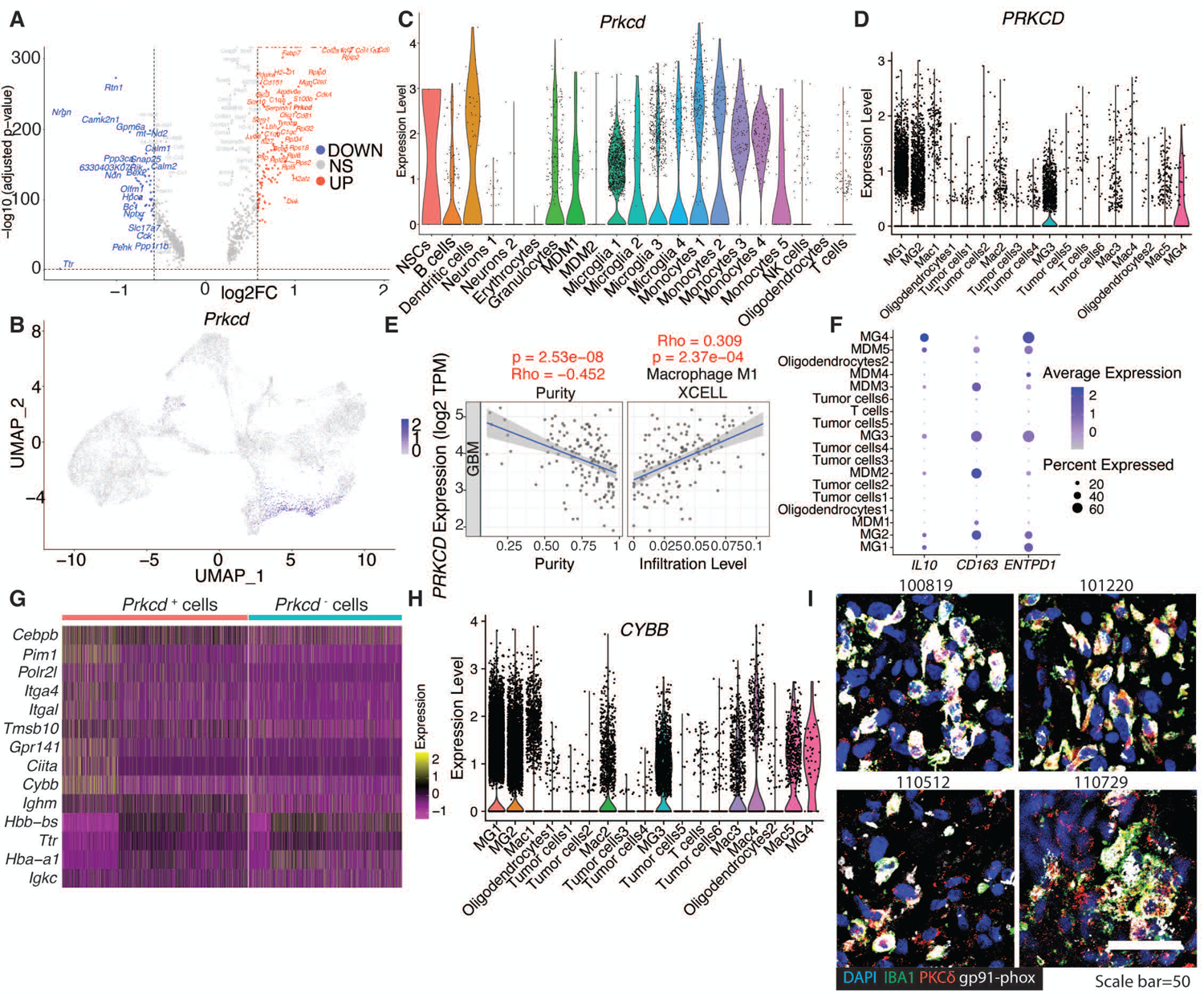
Identification of PKCδ-positive microglia in the GBM microenvironment. A. Volcano plot displaying DEGs between cluster 2 of mouse spatial transcriptomics compared to all other clusters. B. UMAP plot depicting the expression of *Prkcd* in spatial transcriptomics. C. Violin plot showing *Prkcd* expression in scRNA-seq clusters of mouse GBM. D. Violin plot displaying *PRKCD* expression in scRNA-seq clusters of human GBM. E. Correlation analysis between *PRKCD* expression, macrophage infiltration level, and phenotype based on Tumor Immune Estimation Resource (TIMER2.0). F. Dot plot depicting gene expression levels of M2-like macrophage markers in the human tumor microenvironment using human scRNA-seq data. G. Heatmap showing DEGs between *Prkcd*^+^ and *Prkcd*^-^ populations in scRNA-seq of mouse GBM. H. Violin plot illustrating the distribution of *CYBB* in different clusters of scRNA-seq of human GBM. L. Representative IF images of human GBM specimens stained for IBA1, PKCδ, and gp91-phox (CYBB) in four patients.

Next, we unveiled the spatial distribution of immune cell types including subpopulations of monocytes, MDMs, and MG from the scRNA-seq analysis. We also performed gene set enrichment analysis (GSEA) using pathways from the Molecular Signatures DataBase (MSigDB). The hallmark of cancer (HofC), KEGG and Reactome genesets were used to identify enrichment of pathway activity using DEGs from cell types in the scRNA-seq data. Monocyte subsets 1-5 were dispersed throughout the tumor and non-tumor regions, with tendency to accumulate near tumoral margins (**Fig. 2A** and **Supp Fig. 4A**). Notably, monocyte subtype 5 exhibited a unique localization pattern, predominantly infiltrating the tumor core region, unlike the other monocyte subtypes which localized to the tumor periphery. Monocyte subtype 5 was marked by negative enrichment of several immune related pathways from HofC, KEGG and REACTOME. These pathways included the negative enrichment of adaptive and innate immune systems, interferon (IFN) gamma signaling, IFN alpha signaling, MHC class II antigen presentation, interleukin (IL)10 and cytokine signaling within monocyte subset 5 suggesting an immunologically inactive phenotype (**Supp Fig. 5** and **6**). Notably, monocyte subset 5 showed a positive enrichment profile in the cell cycle pathway.

**Figure 5.**
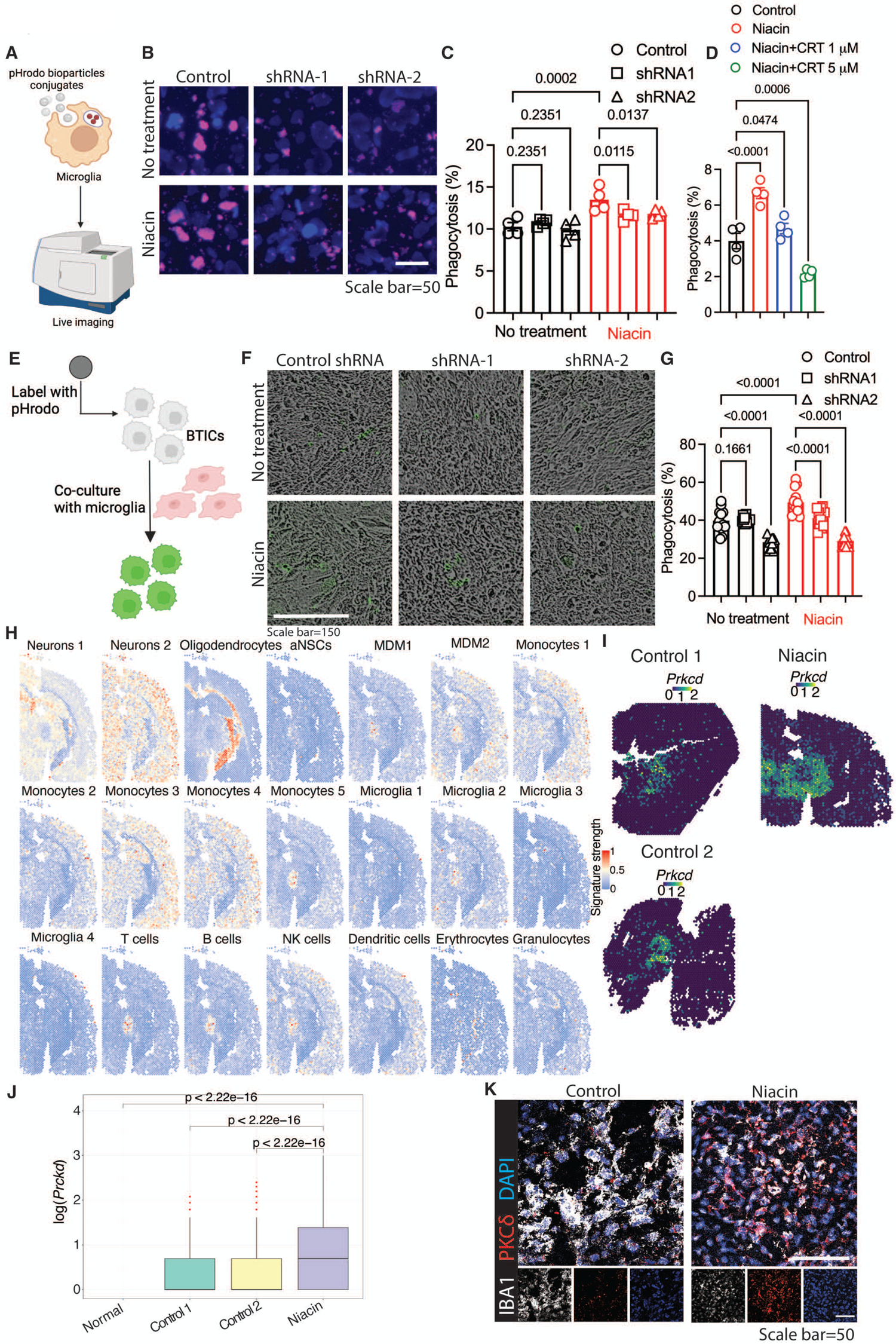
PKCδ in microglia contributes to phagocytosis of GBM cells. A. Schematic overview of the in vitro phagocytosis assay. B. Phagocytosis of pHrodo S. aureus BioParticles by CHME cells with downregulated *PRKCD* expression in the presence or absence of niacin. D. Phagocytosis assay depicting primary human MG stimulated with niacin in the presence or absence of the protein kinase inhibitor CRT 0066101. E. Schematic representation of in vitro phagocytosis of human BTICs conjugated with pHrodo by MG. F. pHrodo-conjugated human BTICs treated with CHME cells in which *PRKCD* expression was downregulated. G. Quantitative analysis of phagocytosis of human BTICs by CHME cells with downregulated *PRKCD* expression. H. Cellular deconvolution of Visium transcriptomics data for tumor mice receiving oral niacin. I. Comparison of *Prkcd* expression levels in niacin-treated mice versus control mice based on Visium transcriptomics. J. Quantification of *Prkcd* expression in Visium transcriptomics. K. IF staining for PKCδ and IBA1 in mouse brains after niacin treatment compared to control. One-way ANOVA with Benjamini multiple-comparison test for comparisons involving two or more treatment groups against the control group. Significant differences in *Prkcd* expression between spatial slides were determined using Wilcoxon rank test (p < 0.05). All data presented as the mean ± SEM (error bars).

**Figure 6.**
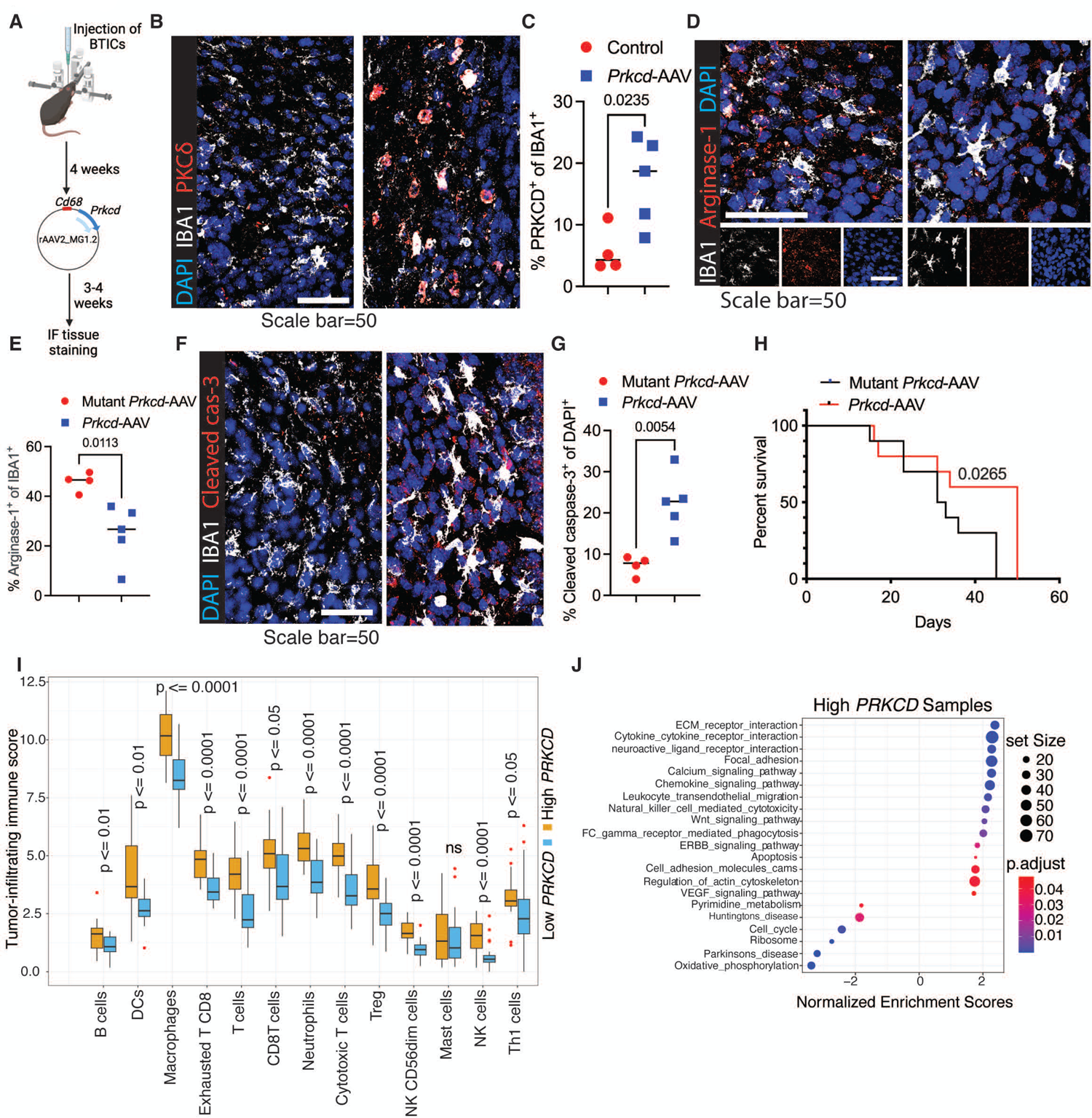
Overexpression of *Prkcd* in microglia and macrophages via adeno-associated virus restrains intracranial tumor growth. A. Schematic of intratumoral injection of AAVs overexpressing *Prkcd* in MG and macrophages after tumor implantation. B. Representative IF images showing expression of PKCδ in IBA1-positive cells after *Prkcd*-AAV injection. C. Quantification of PKCδ-positive cells in the TME of *Prkcd*-AAV-treated mice versus control mice injected with PBS. D. Representative IF images showing expression of arginase-1 in IBA1-positive cells in *Prkcd*-AAV-treated mice compared to mice treated with AAVs expressing a mutant version of *Prkcd* (K376R). E. Quantification of PKCδ-positive cells in the mouse TME. F. Representative IF images of cleaved caspase-3 and IBA1. G. Quantification of cleaved caspase-3 in the TME. H. Kaplan-Meier survival curve of mice treated with *Prkcd*-AAVs and mutant *Prkcd*-AAVs encoding a deficient version of *PRKCD*. I. Tumor-infiltrating immune score in GBM high *PRKCD* verses *PRKCD* low expression of TCGA samples. J. Enrichment pathway analysis of GBM high *PRKCD* verses *PRKCD* low expression of TCGA samples. Significance in panel C, E, and G was determined using a two-tailed, unpaired t-test. Kaplan-Meier survival curves were analyzed for statistical differences between groups using the log-rank Mantel-Cox test. Significant differences in immune scores were determined using Wilcoxon rank test (p < 0.05).

Additionally, MDMs cell populations were localized to the TME in spatial slides (**Fig. 2** and **Supp Fig. 4A**). MDM1 and MDM2 subtypes displayed negative enrichment of HofC, KEGG, and REACTOME pathways including IFN signaling, and cytokine signaling (**Supp Fig. 7**). The pathway results imply that the GBM microenvironment was infiltrated by subtypes of monocyte and MDM populations with immunologically dormant phenotypes, potentially contributing to the unfavorable prognosis of GBMs.

Conversely, immune cell types that showed positive enrichment of immune related pathways indicating immune activity and response were also localized to the TME region. These cell types included monocyte subtypes 1, and 3 (**Fig. 2A** and **Supp.** Fig. 4A). These cell types were positively enriched for HofC, KEGG, and REACTOME pathways including IFN alpha and gamma signaling, cytokine signaling, antigen processing and presentation, IL10 signaling, and JAK and STAT signaling (**Supp Fig. 5** and **6**). Enrichment of MG cell type signatures were limited, but MG subset 2 signatures were present within the tumor area (**Fig. 2A** and **Fig. 4A**).

MG subset 2 showed positive enrichment of HofC, KEGG, and REACTOME pathways including MHC class II antigen presentation, IFN alpha and gamma signaling, cytokine signaling, and TCR signaling (**Supp Fig. 8** and **9**). These results may indicate a dynamic relationship between immune cells that promote immune suppression or immune activity based on the infiltration of specific subtypes of monocytes, MDMs, or MG and thus contribute to cellular heterogeneity within the GBM TME.

In the realm of adaptive immune cells, including T and B cells as well as NK cells, we observed significant accumulation in the tumor core compared to non-malignant areas (**Fig. 2A** and **Supp Fig. 4A**). However, this pattern was not evident for DCs and granulocytes, which exhibited a more uniform distribution across the tissue sections. Interestingly, the tumor areas were devoid of dendritic cells. Altogether, these findings provide a comprehensive understanding of the intricate spatial organization of various non-immune and immune cell populations within the GBM tumor microenvironment.

Given the pivotal roles of MDMs, monocytes, and MG in GBM tumor development and their higher accumulation in the TME, we decided to focus further on these immune cell populations in this study. The tissues from a distinct set of mice implanted with the same tumor cells were subjected to immunofluorescence (IF) staining, utilizing a combination of CD68 and TMEM119 markers to identify MDMs and MG. In alignment with our integrated transcriptomic analysis, our IF data revealed that MDMs predominantly infiltrated near the tumor core (**Fig. 2B** and **Supp Fig. 4B**). Conversely, MG were predominantly located at the outer edges of the tumor core and exhibited dispersed accumulation throughout the entire brain. These findings corroborate with the transcriptomic analyses, reinforcing the distinct spatial distribution of these immune cell populations in the GBM microenvironment.

### Heterogeneous cellular interactions in the GBM microenvironment

In our GBM mouse model, beyond just identifying patterns of cellular localization, we used the CellChat package to decipher the global communication patterns among various cell types found in the GBM TME. Within the deconvoluted cell types from the mouse scRNA-seq analysis, we categorized cell-cell interactions into three distinct patterns. Specifically, cell types including neurons 1, DCs, monocytes 1 and monocytes 2, as well as monocytes 3 and B cells, exhibited pattern 1 of incoming communication (**Fig. 3A**). This pattern suggests that these cells received similar signals from other cell types. On the other hand, MG3, MG4, MG2, MG1, aNSCs, and oligodendrocytes displayed pattern 2 of incoming communication. Finally, pattern 3 was observed in NK cells, granulocytes, T cells, monocytes 5, neurons 2, and MDM2.

We detected significant variations in incoming communication patterns, each intricately linked with distinct signaling networks. Pattern 1 of incoming communications was found to be associated with pathways such as macrophage migration inhibitory factor (MIF), chemokine ligand (CCL), galectin, complement, secreted phosphoprotein 1 (SPP1), protein S (PROS), visfatin, tumor necrosis factor (TNF), vascular endothelial growth factor (VEGF), CD40, IL4, IL2, FASL among others (**Fig. 3B**). Pattern 2 exhibited a unique signature, involving different pathways such as TGFβ, GAS, GRN, colony stimulating factor (CSF), IFN-II, oncostatin M (OSM), B-cell activating factor (BAFF), IL6, leukemia inhibitory factor receptor (LIFR), and insulin signaling networks. Lastly, incoming pattern 3 was associated with CXCL, protease-activated receptors (PARs), IL1, KIT, IL12, epidermal growth factor (EGF), and CD137 networks. These findings provide a comprehensive view of the intricate molecular interactions shaping the communication networks within the GBM microenvironment.

In the analysis of outgoing communication patterns, we observed a shared pattern 1 for exporting signals within the GBM microenvironment among monocyte 4, granulocytes, MG2, DCs, monocyte 3, MG1, monocyte 1, aNSCs, and oligodendrocytes (**Fig. 3C**). Neurons 1, neurons 2, and B cells exhibited pattern 2, while NK cells, T cells, monocytes 2, monocytes 5, MDM, and activated MDM displayed pattern 3 of outgoing communications. These distinct patterns highlight the diverse strategies employed by different cell types in transmitting signals within the complex GBM microenvironment. Pattern 1 of outgoing communications involved a diverse array of signaling molecules, including MIF, galectin, complement, TGFβ, secreted phosphoprotein 1 (SPP1), GRN, CXCL, visfatin, CSF, annexin, TNF among others (**Fig. 3D**).

Signaling in pattern 2 utilized different arrays of pathways such as GAS, midkine (MK), angiopoietin-like protein (ANGPTL), Insulin-like growth factor (IGF), LIFR, and insulin. Lastly, pattern 3 was associated with CCL, PARs, IFN-II, CD40, IL4, IL6, IL2, FASL, and FMS-like tyrosine kinase 3 (FLT3).

Interestingly, immune cell subsets, particularly the monocytes and MDMs were highly concentrated in tumor areas, including monocytes 3, 4, 5, MDM1, MDM2, and faint signatures of MG1, MG2, which received a higher number of signaling networks associated with tumor development, such as galectin, complement, TGFβ, SPP1, TNF, IFN-II, and IL4 (**Fig. 3E**).

Moreover, these infiltrating immune cell populations exported elevated levels of signaling pathways, including CCL, GRN, CXCL, TNF, IL4, IL6, IL2, and IL12, compared to other subsets (**Fig. 3F**). In contrast, MG3 and MG4, and monocytes 1, 2, with less concentrated accumulation in tumor areas, expressed lower levels of signaling strength related to tumor development or immune cell suppression, such as MIF, TNF, IL4, IL6, IFN-II, OSM, galectin, and TGFβ, when compared to other subsets (**Fig. 3E** and **F**). These findings underscore the significance of distinct immune cell populations with diverse spatial localizations in displaying varying interaction profiles within the GBM microenvironment.

In our investigation of the primary contributors to both incoming and outgoing signaling pathways among various cell types, we identified significant roles played by monocytes 3, monocytes 4, MG1, B cells, and monocytes 1 in the incoming signaling pathways (**Fig. 3E**). Regarding outgoing signaling patterns, key contributors included monocytes 3, MG1, neurons 1, and monocytes 4, among others (**Fig. 3F**). In both the incoming and outgoing signaling networks, pathways such as MIF, galectin, complement, TGFβ, and CCL emerged as the strongest contributors for cell-to-cell interaction (**Supp Fig. 10**), highlighting their pivotal roles in regulating cellular communications within the GBM microenvironment.

To validate these findings in human GBM and assess the significance of the top five identified signaling networks from the mouse study, we examined the published scRNA-seq dataset of human GBM. We explored all significant interactions (Ligand-Receptor pairs) associated with MIF, galectin, complement, TGFβ, and CCL within the human GBM microenvironment. Except for MIF, we noted the enrichment of all networks in the human GBM microenvironment.

Specifically, within the galectin network, *LGALS9* was implicated in interactions between various subsets of tumor cells and immune cells (**Fig. 3G**). *LGALS9* expression was observed in MG1, MG2, MDM3, and MDM5, and it played a role in their communication with T cells, MG3, MDM2, and tumor cells 1 and 3. This communication was facilitated through different receptors, including *CD44*, *PTPRC*, and *HAVCR2* on the target cells.

Another significant signaling network identified was the complement pathway. Several subsets of human MG and MDMs, including MG1, MG2, MDM3, and MDM5, expressed C3 complement, facilitating their interaction with MG3 and MDM2 through *ITGAM*_*ITGB2*, *C3AR1*, *ITGAX*_*ITGB2*, and *C3AR1*, respectively (**Fig. 3H**). Similarly, the TGFβ signaling pathway played a significant role in mediating communication between MG1, MG2, MDM3, and MDM5 with MG3, through engaging with *TGFbR1_R2* (**Fig. 3I**). Furthermore, in the CCL network, *CCL3* and *CCL3L3* were primarily produced by MG1, MG2, tumor cells 4, tumor cells 6, oligodendrocytes, and MDM5, facilitating their interaction with MG3 through *CCR1* (**Fig. 3J**). Another chemokine, *CCL5*, was exclusively produced by MDM5 and participated in crosstalk with MG3. In summary, the single-cell and spatial analysis of mouse GBM, in conjunction with human GBM data, revealed critical signaling networks contributing to tumor development within the tumor regions. We also discovered that different immune cells based on their anatomical locations exhibited diverse interaction profiles with others cell types with the possibility to modulate the GBM TME.

### Identification of protein kinase C**δ** (PKC**δ**)-positive myeloid cells in the GBM microenvironment

After identifying significant immune related activity in the GBM microenvironment, we examined individual DEGs in tumor spots more closely for important modulators of immunity. Among the statistically significantly upregulated genes, increased expression of several genes associated with immune regulation was evident in tumoral regions, including *H2-D1*, *C1qa*, *C1qc*, *Ly86*, *Ifi27*, and *Prkcd* (which encodes PKCδ) in the spatial transcriptomics data (**Fig. 4A**). Given the limited information available about the immunological functions of PKCδ in the TME, we conducted a subsequent focused investigation on this gene. Interestingly, spatial transcriptomics data revealed the upregulation of *Prkcd* in both cluster 2 (tumor region) and cluster 7 (tumor adjacent region) in tumor slide (**Fig. 4B** and **Supp Fig. 2H**). These two regions corresponded to areas of the TME where infiltrating immune cells may play a significant role in controlling tumor spread and ultimately influence disease prognosis.

Notably, the scRNA-seq analysis of the mouse GBM samples confirmed elevated *Prkcd* expression in MG, monocytes, and MDMs, particularly in MG2 and monocytes 1 with the highest levels of expression (**Fig. 4C**). Remarkably, based on spatial data, these immune cell type clusters exhibited widespread infiltration across both tumor and non-tumoral areas, showing a notable preference for infiltrating the tumor margin. To validate the cellular distribution of *PRKCD* in human GBM, we analyzed the scRNA-seq data from human GBM samples, revealing predominant expression of *PRKCD* in MG, with higher levels observed in MG1, MG2, and MG3 in that order (**Fig. 4D**).

Using TIMER2.0 (18,19) immune deconvolution analysis and the TCGA GBM RNA data the correlation between *PRKCD* expression and TME purity revealed a significant negative correlation; meanwhile a significant positive correlation was observed between *PRKCD* expression and macrophages (M1-like) displaying an anti-tumor phenotype (**Fig. 4E**). The MG1 cluster of human scRNA-seq GBM also displayed less anti-inflammatory (M2-like) phenotypes compared to other clusters based on lower levels of *IL10*, *CD163* and *ENTPD1* expression (**Fig. 4F**). Intriguingly, examination of outgoing and incoming signaling networks in the human scRNA-seq GBM microenvironment using CellChat indicated that MG1 exhibited fewer immunosuppressive signaling pathways, such as IL10, IL16, IL6, TNF, TGFβ, and galectin (**Supp Fig. 11**). Additionally, as previously outlined, GSEA of MSigDB pathways from the mouse scRNA-seq data showed that monocytes 1 and MG2 subtypes were significant positively enriched for IFN gamma response, IFN alpha response, and inflammatory response (**Supp Fig. 6-9**).

To delve deeper into the phenotype of *Prkcd*-expressing cells, we performed differential gene expression analysis between *Prkcd*^+^ and *Prkcd*^-^ MG, monocytes, and MDMs populations in our mouse scRNA-seq data. Among the top-upregulated genes, we noted high expression of *Cybb* in *Prkcd*^+^ MDMs and MG (**Fig. 4G**). In human scRNA-seq GBM, we similarly observed high expression of *CYBB* in MG1, MG2, and MG3, in that order (**Fig. 4H**). The *CYBB* gene encodes the gp91-phox component of the phagocyte oxidase enzyme complex. This complex plays a crucial role in producing superoxide and other reactive oxygen species, essential for microbial killing (20). To corroborate these findings, we conducted IF staining for PKCδ, gp91-phox, and IBA1 (MG and MDM marker) on GBM tissues from four human patients following surgical resection. In all four patients, we observed co-expression of gp91-phox and PKCδ in MDMs and MG within the tumor microenvironment (**Fig. 4I**). Altogether, these results suggest that MG expressing high levels of *PRKCD* that infiltrate the tumor may possess the ability to phagocytose tumor cells.

### PKC**δ** in microglia contributes to phagocytosis of GBM cells: stimulation by niacin

Given the higher expression of *PRKCD* in MG as indicated by mouse and human scRNA-seq analysis, we conducted further validation experiments focusing on MG. We performed an in vitro phagocytosis experiment utilizing bioparticles labeled with the pHrodo dye which is sensitive to acidic pH environments in endosomes and lysosomes (**Fig. 5A**). pHrodo emits a distinct bright green fluorescence within phagosomes, allowing for the specific detection of intracellular phagocytosis.

In our previous research, we demonstrated that among various agents tested, vitamin B3 or niacin effectively stimulates phagocytosis in macrophages (21,22). We also observed significant elevation of phagocytosis in primary human MG treated with niacin compared with LPS (**Supp Fig. 12A-B**). Therefore, we utilized niacin as a stimulator of MG phagocytosis to investigate the role of PKCδ in the phagocytosis of tumor cells. MG were isolated from mouse brains and exposed to niacin to assess phagocytosis levels. Our results revealed a significant increase in bioparticle phagocytosis upon niacin treatment (**Fig. 5B, C** and **Supp Fig. 12C**). However, this elevation was abrogated when *Prkcd* was downregulated using two distinct shRNAs. To further validate the involvement of PKCδ in phagocytosis, we employed a pharmacological inhibitor of protein kinase C known as CRT 0066101 (23). It was evident that niacin, in the presence of CRT, failed to enhance phagocytosis (**Fig. 5D**), thus supporting the pivotal role of PKCδ in the phagocytosis process.

To better mimic conditions resembling the tumor microenvironment, we utilized patient derived BTICs labeled with pHrodo and co-cultured them with MG isolated from healthy individuals (**Fig. 5E**). We employed in vitro live imaging to monitor the phagocytosis of BTICs. In agreement with the findings from our in vitro mouse models, the downregulation of *PRKCD* in human MG substantially diminished the stimulatory effects of niacin on BTIC phagocytosis (**Fig. 5F** and **G**).

To obtain a comprehensive understanding of niacin’s effects on the tumor microenvironment, we subjected mouse brain tissue isolated from niacin-treated mice to spatial transcriptomics and scRNA-seq. InferCNV analysis identified areas corresponding to tumor regions with chromosomal abnormalities as previously described (**Supp Fig. 13A-D**). After integrating with scRNA-seq data, we deconvoluted the cell types within the tumor microenvironment using our scRNA-seq mouse data and compared the results with those from earlier described tumor mice, collected at the same time point. Our analysis revealed that following niacin treatment, there was a higher level of infiltration of MG2 around the TME compared to untreated mice (**Fig. 5H, Fig. 2A** and **Supp Fig. 4A**). MG subtype 2 was characterized using the same GSEA and MSigDB pathway analysis. As described previously, this cell type was positively enriched for HofC, KEGG, and REACTOME pathways including MHC class II antigen presentation, IFN gamma and alpha signaling, cytokine signaling, and TCR signaling (**Supp Fig. 9**). In comparison to non-treated tumors, MG2 signatures were very limited in the TME. Beyond the cell infiltration, we also observed a significant upregulation of *Prkcd* in niacin-treated mice, especially in the tumor core margin, when compared to controls and normal brain tissue (**Fig. 5I** and **J**). Volcano plots of DEGs showed significant upregulation of *Prkcd* in cluster 2 (tumor region) and cluster 7 (tumor adjacent region) of spatial transcriptomics after niacin treatment (**Supp Fig. 13E** and **F**).

To validate these findings, we subjected additional tissues to IF staining for PKCδ and IBA1, confirming higher PKCδ expression in IBA1-positive cells in niacin-treated mice compared to controls (**Fig. 5K**).

### Overexpression of *Prkcd* in microglia and macrophages via adeno-associated virus restrains intracranial tumor growth

In a proof-of-concept experiment demonstrating the in vivo role of PKCδ in phagocytosis of GBM cells and inhibition of tumor growth, we employed a targeted approach to upregulate *Prkcd* specifically in MG and MDMs. Utilizing a newly developed adeno-associated virus (AAV) named AAV-MG1.2 (24), we aimed to overexpress *Prkcd* in MG and MDMs in mice (**Fig. 6A**). Following treatment of tumor-bearing mice with *Prkcd*-AAV, a substantial increase in PKCδ protein expression was observed in IBA1-positive cells within the TME compared to untreated control mice (**Fig. 6B** and **C**).

In a separate cohort, we administered treatment to tumor-bearing mice using either *Prkcd*-AAVs or AAVs expressing the K376R point mutation (*Prkcd*-KR) in the ATP binding site of the *Prkcd* gene, which exhibits deficient kinase activity (25). Then, brain tissues were subjected to IF staining for arginase, a marker of immunosuppressive myeloid cells. AAV-*Prkcd* treatment led to a significant reduction in double-positive IBA1 and arginase cells (**Fig. 6D** and **E**), indicating a reprogramming of MG toward an anti-tumorigenic profile upon overexpression of wild-type *Prkcd*. As phagocytosis is expected to result in increased apoptotic tumor cells, tissues from mice treated with *Prkcd*-AAV were stained for cleaved caspase-3. Intriguingly, a significant upregulation of cleaved caspase-3 was observed in cells proximal to MG (**Fig. F** and **G**).

Survival analysis further corroborated these findings, demonstrating that *Prkcd* overexpression mediated by AAVs significantly prolonged the survival of mice compared to those treated with AAVs expressing the mutant *Prkcd* (**Fig. 6H**).

### Analyses of *PRKCD* in the TCGA GBM database supports its anti-tumor function

We stratified TCGA GBM bulk RNA-seq data based on *PRKCD* expression and conducted a differential gene expression analysis between *PRKCD* low and high expression groups. Tumors with *PRKCD* high expression exhibited upregulated levels of various ECM molecules, including *COL8A1, COL5A1, COL1A2,* and *LAMA3* (**Supp Fig. 14A** and **B**). Moreover, genes associated with inflammation and immune cell activation, such as *NLRP3*, *NFATC2*, and *PRAM1*, were enriched in *PRKCD* high tumors. Notably, *PRKCD* high tumor samples exhibited significantly higher immune scores using ESTIMATE when compared to low *PRKCD* samples (**Supp Fig. 14C**). Furthermore, using bulk deconvolution analysis of several immune cell types indicated that high *PRKCD* samples had higher levels of immune cells including dendritic cells, macrophages, and various subsets of T cells compared to *PRKCD* low samples (**Fig. 6I**). GSEA and MSigDB analysis using KEGG and REACTOME pathways revealed enrichment of ECM receptor interaction, cytokine receptor interaction, and chemokine signaling pathways in TCGA GBM tumors with *PRKCD* high expression (**Fig. 6J** and **Supp Fig. 14D**). Notably, the apoptosis pathway was also enriched in *PRKCD* high samples, indicating a possible connection to increased apoptosis of tumor cells by PRKCD^+^ MG. Conversely, low *PRKCD* samples showed negative enrichment for apoptosis, phagocytosis, chemokine, and cytokine signaling pathways, and ECM receptor interaction from IPA results of canonical pathway activity (**Supp Fig. 14E** and **F**). Together the results from TCGA GBM analysis were concordant with scRNA-seq analysis indicating that higher levels of *PRKCD* expression can increase anti-tumor response by influencing immune cell infiltration in the TME.

## Discussion

In recent years, breakthroughs in single-cell profiling technologies have transformed our comprehension of tumor heterogeneity, plasticity, and potential therapeutic strategies. Despite these advancements, the significance of the spatial organization within brain tumors has remained largely unexplored. In this study, we pioneered integrative single-cell and spatial RNA sequencing analyses in a GBM mouse model, unraveling the intricate spatial landscape of the brain TME. By pinpointing crucial signaling pathways governing cell to cell interactions and unveiling unique immune cell subsets, we extended our investigation to human GBM through single-cell sequencing and bulk RNA TCGA data, corroborating to our mouse model findings.

This comprehensive approach not only validated our previous discoveries but also led to the identification of a previously unknown MG subset with potent anti-tumor functions within the brain microenvironment.

A significant finding in this study is the elucidation of the spatial landscape of the tumor immune microenvironment in GBM. Distinct cell populations occupied specific regions within the brain, and the spatial architecture of immune cells played a decisive role in shaping their phenotype and functions. This was determined by their microenvironmental context and their interactions with other cells. While a handful of studies have underscored the crucial role of cellular location (17,26), the interaction profiles of immune cells with other cell types have been overlooked.

Notably, this aspect of the tumor immune microenvironment has been elucidated in only a limited number of cancer types (14,27–30). Our study stands out as one of the few shedding light on these nuances in GBM. Different immune cells, residing in diverse spatial locations, displayed varied pathway activity and interaction profiles with neighboring cells in the brain microenvironment. This underscores the significance of cellular localization indicating the interacting partners of immune cells, a crucial aspect for cancer immunotherapy. Rejuvenating immune cell activity requires a deep understanding of their interactome and their communication with the surrounding environment, in addition to their cellular phenotype and localization. Among various signaling networks, we observed that galectin, complement, TGFβ, and CCL are among the pivotal pathways regulating cell-cell interactions in the GBM microenvironment.

Our integrative approach to studying the TME enabled us to unveil previously unidentified MG exhibiting potent anti-tumor functions. These discoveries might have remained elusive if traditional technologies requiring tissue disruption were employed. The identification of MG with anti-tumor phenotypes and functions is crucial (31), as it opens avenues for the development of tools to reprogram suppressed MG toward anti-tumor phenotypes. Through spatial characterization of the GBM microenvironment, we pinpointed PKCδ^+^ MG, present at a very low frequency within the tumor microenvironment. Subsequently, we proposed two tools including vitamin B3 (niacin) and AAV therapy with high potential for translation into clinical applications, aiming to reprogram phenotypes of MG toward an anti-tumor phenotype with high expression of PKCδ. PKCδ, functioning as a cytoplasmic kinase, plays diverse roles by phosphorylating multiple target proteins involved in various cellular processes (32). Its involvement in phagosome-mediated destruction of tumor cells, akin to its role in bacterial infections (33,34), is plausible. The functions of PKCδ in tumor cells remain controversial (35–37). However, in our study we found higher *PRKCD* expression in immune cells, specifically in MDMs, monocytes, and MG, compared to tumor cells within the GBM microenvironment.

Intriguingly, niacin, a *PRKCD* inducer, did not directly impact GBM cell growth in our previous study which showed that it affected myeloid cells (21). Moreover, we employed targeted delivery of the *PRKCD* gene into MDMs and MG using an AAV strategy.

Notably, MG, being the earliest immune cells to combat tumors during the initial phases of tumor growth in the central nervous system (38), offer a strategic window for intervention. Moreover, MG and MDMs are the most prevalent immune cells in brain tumor microenvironment (39). Thus, targeted therapies aimed at reprogramming these cells hold substantial potential to hinder early-stage tumor growth in the CNS.

Reprogramming MG in the brain TME stands out as a promising strategy, especially when contrasted with interventions in later stages characterized by expansive and uncontrollable tumor growth. It is important to note that PKCδ is also expressed by other myeloid cells in GBM, such as MDMs, monocytes and neutrophils. However, the focus on PKCδ in MG is particularly intriguing due to their pivotal roles in brain tumor surveillance. Building upon our previous findings (22) and the current results demonstrating the promotion of PKCδ expression by niacin, along with the targeted in vivo elevation of PKCδ leading to MG reprogramming and tumor growth inhibition, we have launched a phase 1/2a clinical trial to evaluate the safety and outcome of niacin treatment in GBM patients (ClinicalTrials.gov Identifier: NCT04677049).

Collectively, our data indicate that the spatial localization of cells significantly influences the phenotypes, interactome, and functions of immune cells within the GBM microenvironment. We have elucidated the regional landscape of GBM and meticulously mapped its microenvironmental terrain, encompassing the spatial localization of cells and their interactions. This innovative approach to studying the tumor microenvironment has resulted in the identification of a previously unrecognized subset of immune cells, along with the development of two reprogramming tools poised for clinical translation. Altogether, these approaches have proven effective in resolving tumor heterogeneity and generating a better understanding of GBM and its immunosuppressive microenvironment.

## Methods

### Animal experimental models

In this study, all mice utilized were of C57BL/6 background and were sourced from Charles River as 6- to 8-week-old female wild-type mice. Ethical approval for all experiments involving animals was obtained from the Animal Care Committee at the University of Calgary, following the guidelines set forth by the Canadian Council of Animal Care.

### Microglia isolation from human

MG were extracted from mixed preparations derived from brain tissue of human fetuses aged between 18-22 weeks gestation, following established protocol (40). The purity of the MG obtained exceeded 95%. The use of these specimens was approved by the University of Calgary Research Ethics Board. The MG were diluted and fed using a medium composed of MEM, L-glutamine, sodium pyruvate, 1% penicillin/streptomycin, 10% dextrose and 10% fetal bovine serum (all culture constituents were from Invitrogen).

### Microglia isolation from mice

Primary mouse MG, constituting over 95% pure non-adherent cells, were cultured from neonatal brains using methods described in a previous publication (41).

### Microglia cell line

CHME human MG cell line was used for in vitro validation experiments such as down regulation of *PRKCD* in MG. This cell line was cultured under the same condition as primary human MG.

### Human GBM tissues

For immunohistochemical analyses, human GBM tissues (n = 4) were obtained in accordance with protocols approved by Indiana University’s Institutional Review Board. Fresh tissues were procured from patients through surgical procedures, immediately snap-frozen, and stored at −80°C. Subsequently, the tissues were fixed in formalin and processed for histological examination, leading to the creation of paraffin-embedded blocks. Hematoxylin and eosin (H&E)–stained slides were meticulously examined by a neuropathologist. Written informed consent was obtained from all patients prior to the collection of tumor biopsies. The information of patients related to these specimens was described previously (5,42). We thank Dr. Chunhai Hao, Indiana University, for providing the GBM samples.

### Generation of GBM patient derived BTICs

Human BTIC lines were derived from resected tissues of GBM patients and cultured following established protocols (21,43,44). These lines were sequentially cultured, preserved, and authenticated within the University of Calgary BTIC Core. Previously reported identification of two human BTIC lines (BT048 and BT073) was utilized in this study (5,42). Short tandem repeat analysis was conducted for cell line authentication, and routine testing for Mycoplasma contamination was performed using the Venor GeM Mycoplasma Detection Kit (Sigma, MP0025).

### Live cell imaging of in vitro phagocytosis

MG were seeded in a 96-well black-bottom plate at a density of 30,000 cells per well in complete medium and incubated for 24 hours at 37°C with 5% CO2. Following incubation, the medium was replaced with DMEM containing 1% FBS. After 1 hour of incubation, niacin (100 μM) was introduced to the cells, and they were further incubated for 12 hours at 37°C with 5% CO2.

Subsequently, the medium was aspirated, and pHrodo S. aureus BioParticles (ThermoFisher, P35361) or BTICs labeled with pHrodo Red, succinimidyl ester (ThermoFisher, P36600) were added to the cells in live cell imaging solution. To stain cell nuclei, NucBlue Live ReadyProbes Reagent (two drops per mL; Thermo Fisher, R37605) was also added. The cells were then observed using an ImageXpress Micro Cellular Imaging and Analysis System (Molecular Devices) for one hour. For continuous monitoring of phagocytosis, the Incucyte microscopy system was employed. The percentage of NucBlue-positive cells that displayed pHrodo positivity was quantified using MetaXpress or Incucyte software.

### Intracranial tumor implantation

After dissociation of spheres, 25000-50000 viable cells from firefly luciferase–expressing or wild-type BTIC0309 were resuspended in 2 μL of PBS and stereotaxically implanted into the right striatum of mice, following previously described methods (45). In the niacin experiments, starting seven days after implantation, mice were treated with niacin at a dose of 100 mg/kg daily via intraperitoneal injection until the termination of the experiment. Control mice were injected with an identical vehicle (saline) solution.

### Preparation of libraries for spatial transcriptomics

The spatial transcriptomics analysis was conducted using the Visium Spatial Gene Expression platform by 10x Genomics. Brain tissues, collected 40 days after tumor implantation, were frozen and sectioned into 10-μm slices, following the manufacturer’s guidelines. Sections were placed on Visium slides and processed as per the user guide. Tissue optimization studies delineated an optimal permeabilization time of 12 mins. Brightfield images distinguishing tumor and non-tumor regions were captured using the EVOS FL Auto Imaging System. The subsequent library preparation steps were performed as per the user guide (Visium Spatial Gene Expression Reagent Kit; CG000239). The libraries were then sequenced on an Illumina NovaSeq 6000 system, employing a NovaSeq 200 Cycle S1 flow cell and specific read cycles for sequencing.

### Data processing and batch correction of spatial transcriptomics

Spatial transcriptomics data was imported (images, counts, scale factors, and tissue position files) using Seurat and the CleanSpot R packages. Four tissue slides/samples were merged containing one normal tissue slide with no tumor, one tissue slide with tumor and niacin treatment, and two tissue slides with tumor and no niacin treatment. Initial data processing removed lowly expressed genes and spots with low feature counts; no spots were removed with mitochondrial contamination because removal of these spots compromised the overall number of spots available for analysis.

The merged spatial Seurat object was normalized (SCTransform, PCA components = 30), scaled, and then spots were clustered (res = 0.5) as per basic Seurat processing protocols. Next, spatial slides were checked for sample-based batch effects, and then these effects were removed using the Harmony R package. After batch correction, spots clustered based on biological gene expression and not by tissue slide/ samples. After initial data processing there were 12,588 spots ready for downstream analysis. After default Seurat clustering, differential expression analysis was performed between Seurat clusters using the FindAllMarkers function, thresholds for DEGs were set to log2fold change +/- 0.58 and p-adjusted value < 0.05.

### InferCNV – Cancer/Tumor Spot Identification

Copy-Number Variation (CNV) events were used to identify tumor cells within spots of spatial slides. The InferCNV pipeline was run as per default, but importantly noting we used the Seurat clusters designated as “cancer spots – cluster 2” and “normal spots – all other clusters” to compare and identify CNV events in cancer vs normal spots. Tumor cells/spots were identified using the top duplication event identified, tumor spot identification overlapped nicely with visual images of tumor location on the tissue slide. Differential expression analysis was performed between identified cancer/ tumor spots and adjacent normal spots to validate InferCNV calls by identifying cancer-associated upregulated genes within cancer/ tumor spots.

### Spot deconvolution in spatial transcriptomics

Spatial deconvolution was performed with the CARD R package to assign cell type identity to spots by integrating our in-house mouse GBM single-cell data as a reference. CARD uses autoregressive-based deconvolution to combine cell type specific expression information from scRNA-seq data with correlation in cell type composition across tissue locations. Each spot on the tissue slide was assigned a proportionality score of multiple cell types detected within each spot. To validate and confirm CARD results we also performed deconvolution using the SingleR to identify concordant calls between the two methods.

### Analysis of public scRNA-seq datasets

The counts matrix of Richard et al (4) was downloaded and converted into a Seurat object using the Seurat v3 package in R (46). Initial data filtering involved excluding cells with less than 50 genes and a percentage of mitochondrial genes exceeding 15%. Following this, data were integrated and normalized using the SCTransform wrapper in Seurat. Principal component analysis (PCA) reduction was employed, considering 15 significant PCA dimensions. Clusters were identified using the FindNeighbours and FindClusters functions, with clustering executed at a resolution of 0.5. Manual annotation of clusters was carried out based on the expression patterns of lineage-specific signature genes. Cell clusters were visualized using the RunTSNE function with PCA reduction. Differential expression genes (DEGs) for individual clusters (versus all cells in other clusters) were identified using the FindMarkers function, including only DEGs with a statistically significant P value of 0.05 for subsequent analysis. To visually represent the results, the DoHeatmap functionality in Seurat was utilized to generate heat maps of DEGs of interest for each cluster. Dot plots illustrating the average expression of hallmark genes and genes of interest, expressed as a percentage in each cluster, were generated using the DotPlot function. Additionally, the FeaturePlot and VlnPlot functions were employed to create graphs depicting gene expression in specific cell clusters or sample groups.

### Confocal immunofluorescence microscopy

We analyzed samples from GBM patients fixed in formalin and embedded in paraffin, as well as frozen mouse sections, following established protocols (5,42). Following antibodies were used for tissue staining: PKCδ (Abcam, ab182126, 1:200 dilution), IBA1 (ThermoFisher, PA5-18039, 1:100 dilution), Arginase (Cell Signaling, 93668, 1:50 dilution), Cleaved Caspase-3 (Cell Signaling, 9661, 1:500 dilution), TMEM119 (Abcam, ab209064, 1:400 dilution), CD68 (Biolegend, 137001, 1:500 dilution) and gp91-phox (Novus Biologicals, NBP1-41012, 1:100 dilution). After primary antibody incubation, samples were treated with corresponding fluorophore-conjugated secondary antibodies (1:500; Jackson ImmunoResearch Laboratories or Thermo Fisher Scientific) and 4′,6-diamidino-2-phenylindole (DAPI; 1:1000). Laser confocal IF imaging was performed at room temperature using the Leica TCS SP8 microscope with consistent settings across samples, ensuring optimal contrast and minimal saturation. Each sample generated three to four fields of view (FOV). Image acquisition was done using Leica Application Suite X, and subsequent picture thresholding and three-dimensional (3D) rendering were conducted with Imaris software (v9.9.1 Bitplane).

### Confocal image analysis

Z-stacks were imported into Imaris in their original format. Cells expressing specific markers were identified in Imaris utilizing the surface tool. Background subtraction was applied to distinguish cells from the background environment. Accurate quantification was achieved through fluorescence thresholding and masking of undesired immuno-labeling. To identify positive signals, a consistent color brightness threshold was established using a value determined from tissues stained with negative secondary antibodies. The same threshold values for color brightness, along with size and circularity settings, were uniformly applied across all samples within each experimental set. Double positive cells were quantified using the parameter of the shortest distance between surfaces. Statistical data were exported to Excel for further analysis.

Representative images were generated by merging and displaying the maximum-intensity projection of each channel/marker in a z-stack using pseudocolors in ImageJ. Adjustments to brightness and contrast were made consistently across samples to enhance image clarity.

Lentiviral vectors containing validated shRNA sequences targeting expression of human *PRKCD* (accession NM_006254) or a non-target control were obtained from Sigma (catalog numbers TRCN0000010193, TRCN0000195408, SHC002). Lentivirus production was performed using 293FT cells grown to 90% confluency on five, 15 cm^2^ cell culture plates, by cotransfecting 112.5 μg lentiviral vector, 73 μg psPAX2 and 39.5 μg pMD2.G [gifts from Didier Trono (Addgene, plasmid # 12260) using PEIpro transfection reagent (Polyplus). The cell culture media was collected 48 hours post-transfection, centrifuged at 500xg for 5 min and 0.45 um filtered. The filtered media was underlaid with a 10% sucrose cushion and centrifuged at 12,000 xg for 4 hours at 4°C (47). The resulting pellet was resuspended in sterile 1XDPBS and frozen at −80°C as 20 µL aliquots. The lentivirus titer determined using the qPCR lentivirus titer kit was >10^9^ IU/mL (Applied Biological Materials).

### Plasmid construction

pAAV-CD68-hM4D(Gi)-mCherry, with the monocyte-specific promoter CD68 and was a gift from B. Roth (Addgene plasmid no. 75033(48)), was digested with SalI and HindIII to excise the hMD(Gi)-mCherry sequence. The cDNA sequence corresponding to mouse *Prkcd* (accession AF251036) was amplified from a mouse brain cDNA library and subcloned to generate pAAV-CD68-*Prkcd* or pAAV-CD68-*Prkcd*-KR (K376R mutant with ATP binding deficiency (25)) using the NEBuilder Hifi DNA assembly kit. All DNA constructs were verified by Sanger DNA sequencing.

### AAV production for in vivo overexpression of *PRKCD* in microglia and macrophages

AAV viral vectors containing the MG1.2 capsid were generated using previous methods (49). Briefly, 293FT cells were grown to ∼90% confluency in Corning Hyperflasks and co-transfected with 129 μg of pHELPER (Agilent), 238 μg of Rep-Cap plasmid encoding MG1.2, a gift from Minmin Luo (24) (Addgene plasmid # 184541) and 64.6 μg pAAV-*CD68-Prkcd* or pAAV-*CD68-Prkcd*-KR using the PEIpro transfection reagent (Polyplus). AAVs were precipitated from medium collected after 3 d and 5 d using a solution of 40% PEG/2.5 M sodium chloride and pooled with cells harvested after 5 d in buffer containing 500 mM sodium chloride, 40 mM Tris Base and 10 mM magnesium chloride. The lysate was incubated with 100 U ml−1 salt-active nuclease (Arcticzymes) at 37 °C for 1 h and then centrifuged at 2,000*g* for 15 min. AAVs were purified from the resulting lysate using an iodixanol step gradient containing 15, 25, 40 and 60% iodixanol in Optiseal tubes (Beckman) followed by centrifugation at 350,000*g* using a Type 70 Ti ultracentrifuge rotor (Beckman). Following centrifugation, the AAVs were collected from the 40% layer using a 10-cc syringe and a 16-gauge needle, diluted in 1× PBS containing 0.001% pluronic F68 (Gibco) and filtered using a 0.2-μm syringe filter. The AAVs were concentrated and buffer exchanged by five rounds of centrifugation using Amicon Ultra-15 100-kDa-cutoff centrifugal filter units (Millipore). The titer was determined using the qPCR Adeno-Associated Virus Titration kit (Applied Biological Materials) and the purity was verified by SDS–PAGE and total protein staining using InstantBlue reagent (Expedeon).

### In vivo overexpression of *Prkcd* in myeloid cells with AAVs

To overexpress *Prkcd* in the brain of tumor-bearing mice, a solution of 5 μl saline containing 2.5 × 10^10 genome copies of AAV-MG1.2, designed with MG-targeting potential, was intratumorally injected. The AAV-MG1.2 construct was engineered to overexpress *Prkcd* under the control of the CD68 promoter. This procedure was performed 2 weeks after tumor injection. Control mice were injected with 5 μl of saline containing 2.5 × 10^10 genome copies of AAV-MG1.2 with a CD68 promoter designed to express a mutant version of *Prkcd* (*Prkcd*-KR).

### Single cell isolation for scRNA-seq

Mice were euthanized four weeks post-tumor implantation, and brains were isolated. To enrich for tumor tissue, brains were hemisected, and a 2 mm section around the tumor injection site was collected into Hank’s Balanced Salt Solution (HBSS, Gibco). Four brain tissues were pooled together per sample to ensure an adequate number of cells for downstream scRNA-seq analysis (Control: 5 mice, one replicates; niacin injection: 10 mice, two replicates). Subsequently, brain tissues were homogenized using a Dounce homogenizer and filtered through a 40-µm cell strainer. The cell suspension underwent Percol gradient separation to remove myelin debris. After further filtration through a 35-µm cell strainer, cells were resuspended in PBS containing 1% heat-inactivated FBS (Sigma) and 0.1 mM EDTA.

### scRNA-seq sample and library preparation

Once a single cell suspension was generated as described above, cells were labelled with SYTOX™ Green Dead Cell Stain (ThermoFisher, S34860) to discriminate between live/dead cells. Live cells were sorted on FACSAria III (BD) using a 100-µm nozzle. scRNA gene expression libraries were prepared using the Chromium Next GEM single cell 3’ reagent kits v3.1 from 10x Genomics. Appropriate volume of cells, as determined from the user guide for recovery of 3000 cells, was loaded on the Chromium single cell controller chip. Post cDNA amplification QC and quantification as well as library construction QC was done using an Agilent Bioanalyzer high sensitivity DNA chip for use with the Agilent 2100 Bioanalyzer. For sequencing, all libraries were pooled and loaded at 300pM on the NovaSeq Ilumina system using the NovaSeq S1 flowcell. A 28 base-pair Read 1 was used to sequence the cell barcode and UMI, an 8 bp i7 index read was used to sequence the sample index and a 91bp Read 2 was used to sequence the transcript using paired-end sequencing.

### Single-cell RNAseq Data Processing

Sequencing depth obtained ranged from approximately 95,000-116,000 reads/cell. BCL files obtained were processed using the Cell Ranger version 3.1 pipeline. Briefly, the fastq files were processed with the cellranger count pipeline which uses STAR to align the reads to the mm10 mouse reference transcriptome. The cellranger aggregate pipeline was run to generate an expression matrix with all combined libraries with normalization setting set to ‘None’. The Seurat R package was used to process single cell counts, cells with >200 nfeatures were removed, as well as cells with <20% mitochondrial genes. In total, after filtering, there was 5440 cells ready for downstream analysis. Unfortunately, the tumor cells were lost during sample processing and not available for analysis. The Seurat R package was used to perform data normalization, scaling, and clustering (res = 0.5) of cells, resulting in 17 unique clusters. Differential expression analysis was performed with the FindAllMarkers functions grouping by Seurat clusters to identify top upregulated genes per cluster.

### Cell type deconvolution of scRNA-seq data

scRNA-seq cell-type deconvolution analysis was performed using CellDex and SingleR R packages with the mouse bulk RNA-seq cell type database used as a reference from CellDex. For this analysis we were unable to identify a suitable “already labelled” single-cell mouse brain/ GBM dataset to deconvolute with, thus we settled for a bulk RNA-seq based approach. Cell type labels determined by SingleR were validated using known marker genes for identified cell types. After performing initial cell type deconvolution, we noticed that overall cell type assignments encompassed multiple Seurat clusters. From this observation we performed a refined reclustering analysis to identify subpopulations of cell types based on underlying Seurat clusters. We used the Seurat clusters, DEGs between clusters, pseudotime/ trajectory analysis using the moncole3 R package, and inferred biological pathway activity (GSEA and MSigDB – hallmark of cancer gene sets, GO pathways, KEGG, and REACTOME using the ClusterProfiler R package). From this analysis, we identified 5 subpopulations of monocytes (monocytes 1 and 3 were similar – Group 1, monocytes 2 and 4 were similar – Group 2, and monocyte 5 – Group 3). From the monocyte trajectory analysis, there were three distinct trajectories identified, along pseudo time, Group 1 was timepoint 2, Group 2 were timepoint 3, and Group 3 were timepoint 1. There were two populations of macrophages identified by SingleR, but these cell types did not contain any subpopulations after re-clustering analysis and were labelled MDM1 and MDM2. Using the same methods, we identified 4 subpopulations of MG. Based on DEGs and trajectory analysis, each subpopulation was unique with some similarity between MG 1 and 2 subtypes. MG1 was pseudotime point 2, MG2 was timepoint 3, MG3 was timepoint 1, and MG4 was timepoint 1.

### CellChat – Cell to cell interaction analysis

The mouse GBM scRNA-seq dataset was used as input for a CellChat analysis, and cell type identities were set to SingleR deconvolution/ reclustering analysis results. The mouse proportion of the cell-to-cell interaction database was used, along with secreted ligand, ECM receptors, and cell-cell contact databases. CellChat identified differentially expressed ligand-receptor genes between cell types, and identified significant pathways related to cell-to-cell interactions. The basic CellChat pipeline was used to identify interactions between cell types and incoming/ outgoing interactions was determined using non-negative matrix factorization (NMF).

### Bulk RNA-seq TCGA GBM data download and processing

Glioblastoma (GBM) mRNA-seq expected counts and normalized transcripts per million (TPM) counts were downloaded from the UCSCXena browser under the TCGA Hub. Expected counts were used for DESeq2 differential expression analysis and TPM normalized counts were used for all other downstream applications, including immune cell deconvolution analysis. For DESeq2 differential expression analysis, GBM mRNA-seq counts were used to examine *PRKCD* expression. GBM tumor samples were split into four quantiles of expression based on *PRKCD*, as quantile 1 = low expression, and 4 = high expression. The GBM samples were filtered for expression quantiles 1 and 4, and then DESeq2 differential expression analysis was performed between these two expression groups with default settings. Differentially expressed gene thresholds were set at a log2fold change +/- 0.58 and p-adjusted value < 0.05. DEGs were visualized using volcano plots and heatmaps generated using ggplot2 and pheatmap functions.

### Immune Contexture Analysis

The immune cell contexture of GBM high *PRKCD* verses *PRKCD* low expression of TCGA samples were examined using two methods. Firstly, the ESTIMATE algorithm was used to examine immune scores indicating overall levels of immune cell infiltration between the two expression groups, which also included ESTIMATE stromal and purity scores. Secondly, a paper published using immune cell markers validated by flow cytometry, immunohistochemistry, and know cell type marker databases was used to deconvolute and assign tumor-infiltrating immune (TIL) scores per TCGA sample including GBM samples. *PRKCD* expression groups (high = 4 vs low = 1) were compared for differences in tumor-infiltrating immune scores using Wilcoxon T-test (p-adj value < 0.05). The TIL score dataset came from Danaher et al., 2017 (50).

### Qiagen Ingenuity Pathway Analysis of single cell and TCGA data

Differential expression results from scRNA-seq and TCGA analysis were used to run an IPA core analysis and examine canonical pathway and upstream regulator activity between cell types (scRNA-seq) and expression groups (*PRKCD* High vs Low – TCGA data). Differential expression results from Seurat FindAllMarkers were used as input for IPA using previously defined log2 foldchange and p-adjusted value for scRNA-seq data. Two differential expression analysis were performed, one with DEGs in high *PRKCD* expression samples, and DEGs in low *PRKCD* expression samples in TCGA data. Two separate IPA core analysis were run on the two separate gene sets for TCGA. IPA results were downloaded and visualized using the ggplot2 package.

### Pathway Analysis using ClusterProfiler R Package to perform GSEA of MSigDB Pathways

Secondary pathway analysis was performed using the Clusterprofiler R package, to perform geneset enrichment analysis (GSEA) to identify enriched geneset/ pathways from the Mutational Signatures Database (MSigDB). This database contains gene sets from various sources and for this analysis included the hallmarks of cancer gene sets, KEGG, REACTOME, and Gene Ontology (GO) pathways. The pathway analysis was used DEGs from DEAs to identify the enrichment of pathway activity across scRNA-seq and bulk RNA-seq data. This method was used to identify pathway activity in immune cell types in mouse and human scRNA-seq data using DEGs of each cell type. This method was also used for TCGA GBM RNA data for DEGs from *PRKCD* high vs low samples.

## Data availability

Raw reads for the scRNA-seq, and spatial transcriptomics will be deposited in the Gene Expression Omnibus (GEO) which will be available upon acceptance of the manuscript. All other data are available upon request.

## Code availability

Mouse scRNA-seq and spatial transcriptomics datasets discussed in this paper can be provide as per reasonable request from the authors.

## Statistics and Reproducibility

Data were organized using Microsoft Excel, and graphs were created using GraphPad Prism 10.1.0. Each graph illustrates individual data points, with each point representing either an individual mouse (for in vivo experiments) or a distinct well in a 96-well plate (for in vitro experiments). The data points are accompanied by the mean ± SEM. The sample size was determined based on previous published studies (51,52), taking into account factors such as experimental cost, feasibility, and the availability of sex- and age-matched mice. Specific sample sizes are provided in the figure legends, and only one measurement was taken per sample. Littermate mice were randomly assigned to each experimental condition and treatment. Statistical analysis was performed using one-way ANOVA with Benjamini multiple-comparison test for comparisons involving two or more treatment groups against the control group. For data involving only two groups, significance was determined using a two-tailed, unpaired t-test. Blinding procedures were not implemented during the experiments.

## Authors Contributions

R. Mirzaei: Conceptualization, data curation, software, analysis, validation, visualization, methodology, writing–original draft, writing–review and editing. R. McNeil: Data curation, analysis, visualization, methodology, writing–review and editing. B. Wong: Data curation, writing–review and editing. C. D’Mello: Software, analysis, Single-cell gene expression and spatial library preparation, writing–review and editing. S. Sarkar: Methodology, writing–review and editing. F. Visser: Methodology, writing–review and editing. C. Poon: Writing–review and editing. P. Bose: Supervision, software, writing–review and editing. V.W. Yong: Conceptualization, resources, supervision, funding acquisition, validation, project administration, writing–review and editing.

## Acknowledgments

We thank members of the Yong laboratory for their thoughtful feedback on the study. The authors acknowledge the support and access to the Leica TCS SP8, ImageXpress Micro XLS High-Content Analysis System, and image analysis platforms provided by the Hotchkiss Brain Institute Advanced Microscopy Platform (HBI AMP) and the Cumming School of Medicine. The authors are thankful for the assistance and utilization of the IncuCyte Live-Cell Analysis System at the Snyder Institute’s Live Cell Imaging Resource laboratory at the University of Calgary. The flow cytometry core and the UCDNA sequencing at the University of Calgary were also acknowledged for their support. R. Mirzaei received fellowship support from the University of Calgary’s Eyes High program. Financial support for this study was provided by grants from the Canadian Institutes of Health Research and the Canadian Cancer Society.

---

**Supplementary Figure 1.**
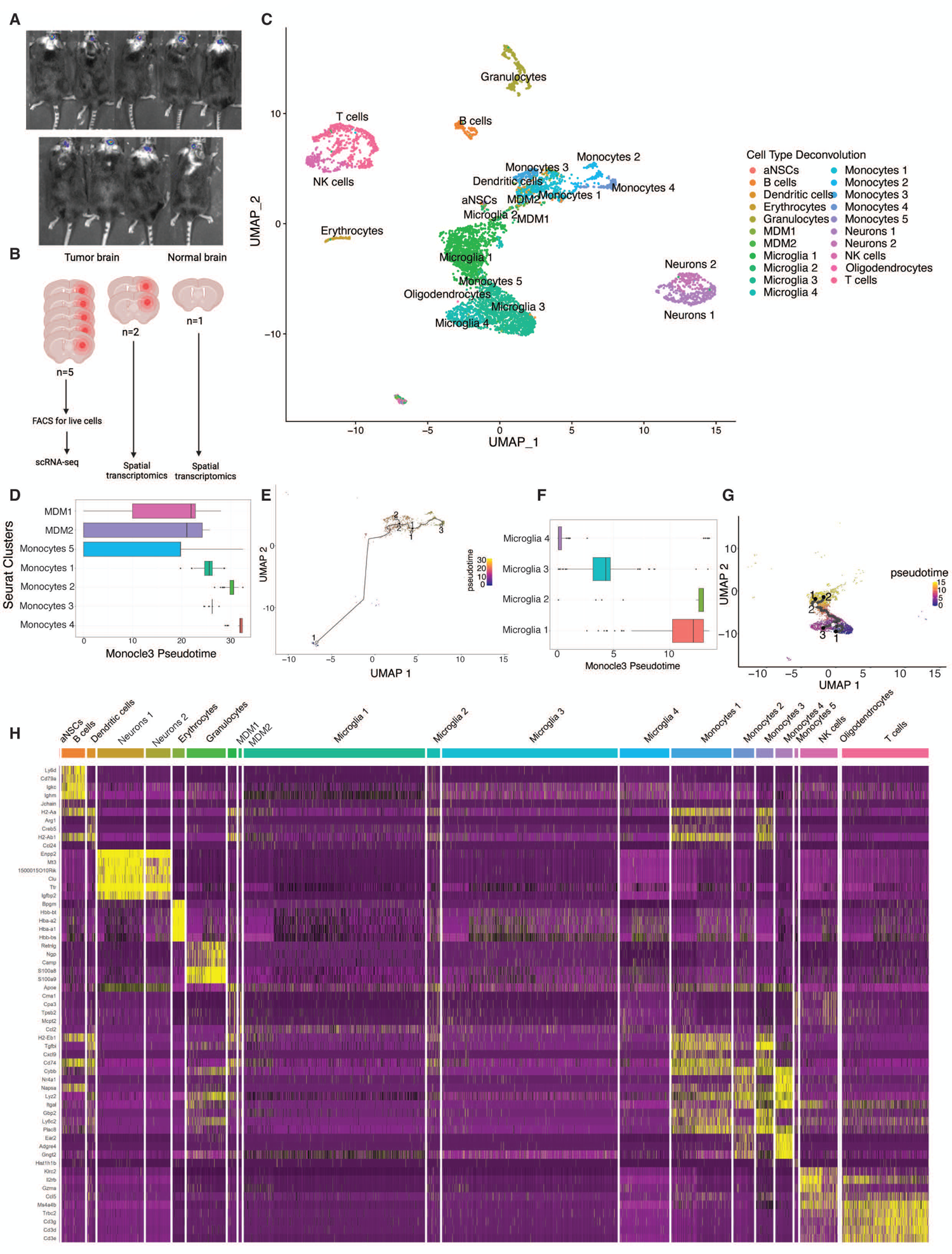
Single-cell spatial analysis of GBM microenvironment. A. Bioluminescence imaging of mice implanted with BTICs before harvesting tissues for scRNA-seq. B. Schematic showing number of mice subjected to scRNA-seq and spatial transcriptomics. C. UMAP plots displaying Seurat clusters identified through scRNA-seq. D. Pseudotime bar plot of different monocyte and MDM sub-clusters showing differences in trajectories. E. UMAP of Pseudotime analysis for monocytes and MDMs showing different trajectories. F. Pseudotime bar plot of different MG sub clusters showing differences in trajectories. G. UMAP of Pseudotime analysis for MG showing differences in trajectories. H. Heatmap of DEGs of scRNA-seq clusters.

**Supplementary Figure 2.**
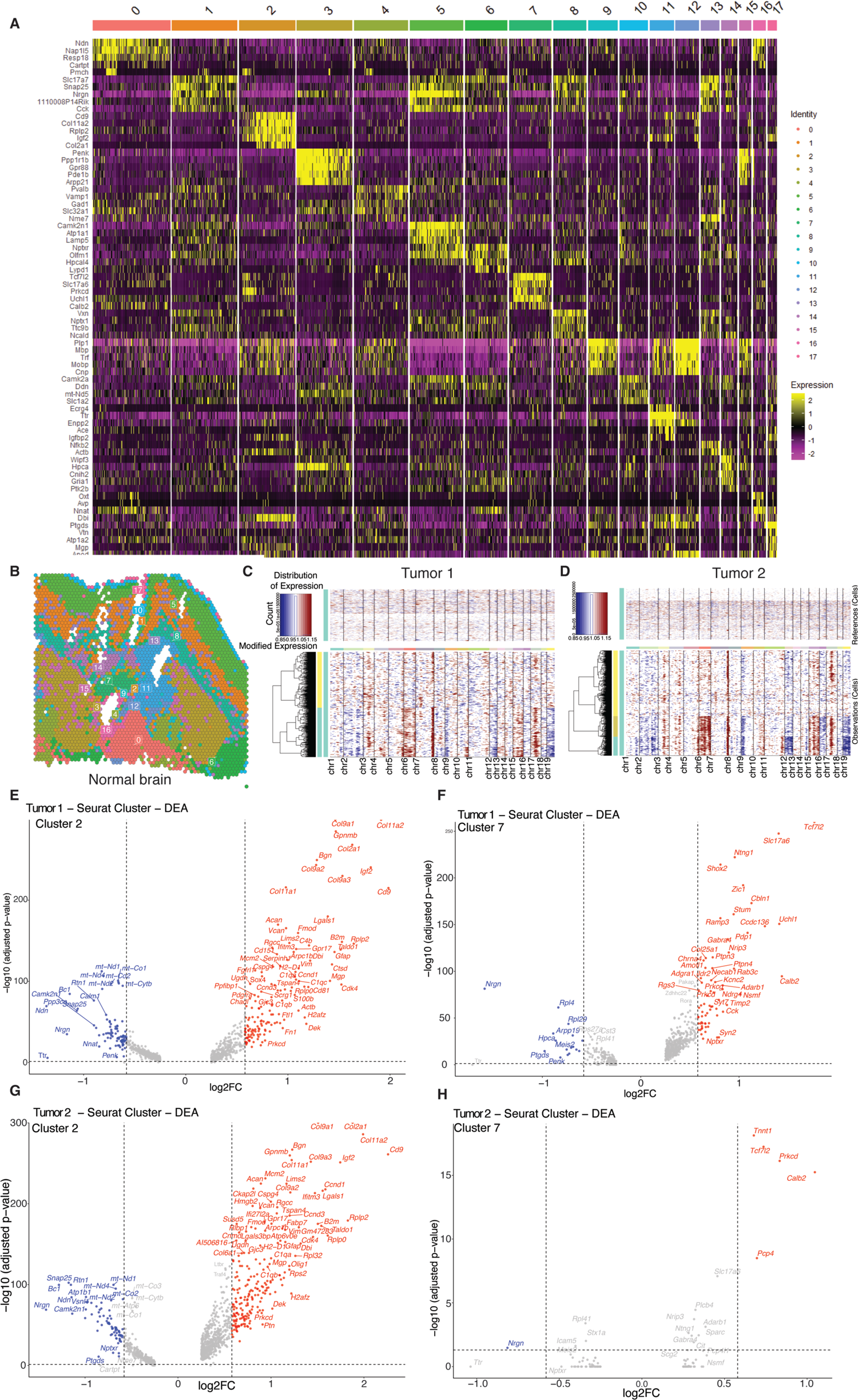
Spatial transcriptomics of mouse GBM. A. Heatmap of DEGs in spatial clusters. B. Seurat spatial clusters overlaid on tissue image of normal brain. C and D. InferCNV plots of two tumor mice subjected to spatial transcriptomics. E-H. Volcano plots of DEGs in cluster 2 (tumor region) and cluster 7 (tumor adjacent region) for spatial transcriptomics of mouse GBM.

**Supplementary Figure 3.**
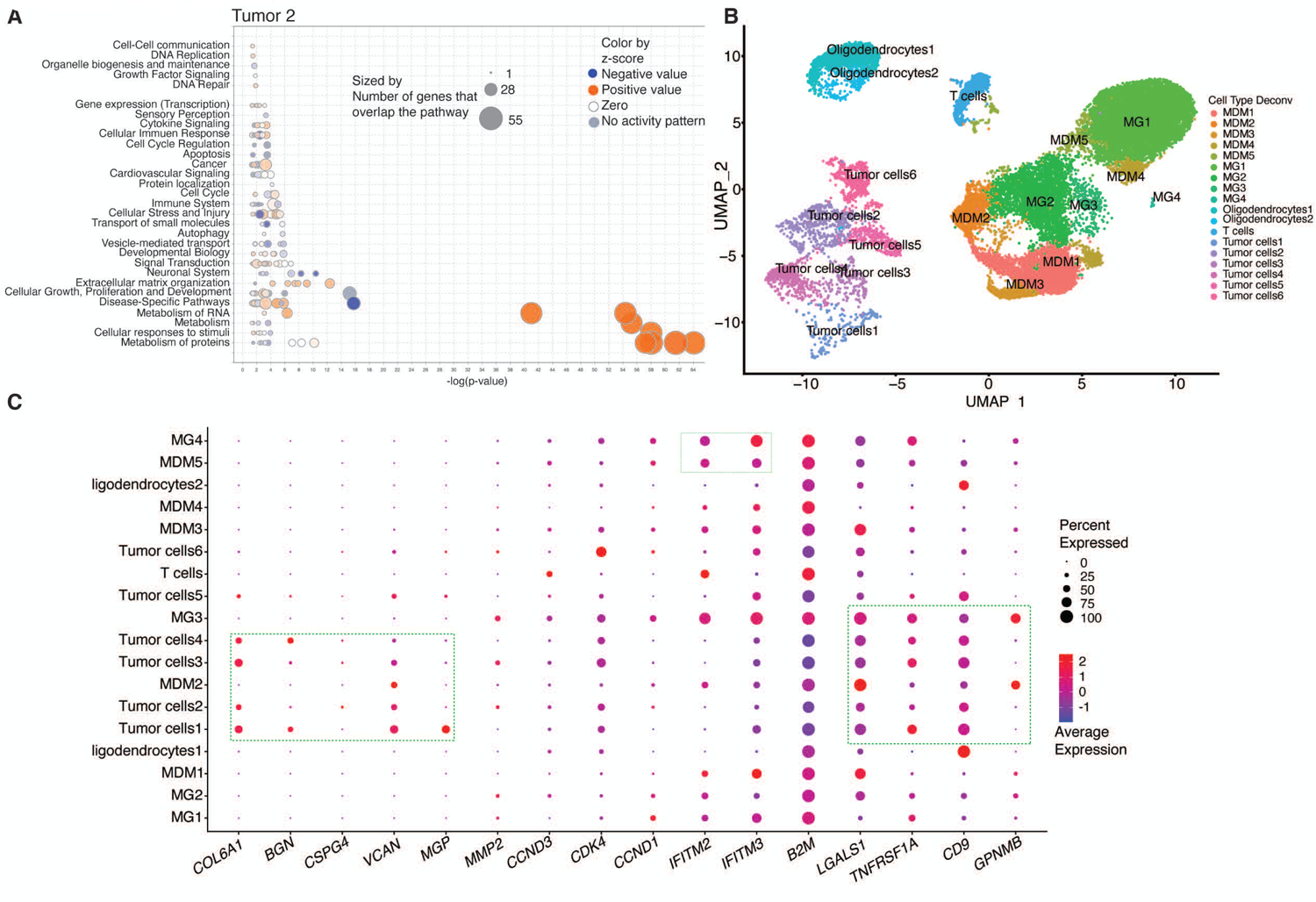
Transcriptomic profile of GBM microenvironment. A. Canonical pathways enriched in DEGs between malignant and non-malignant regions in mouse tumor 2. B. UMAP plots displaying Seurat clusters identified through scRNA-seq of human GBM. C. Dotplot depicting gene expression levels in the human tumor microenvironment using human scRNA-seq data.

**Supplementary Figure 4.**
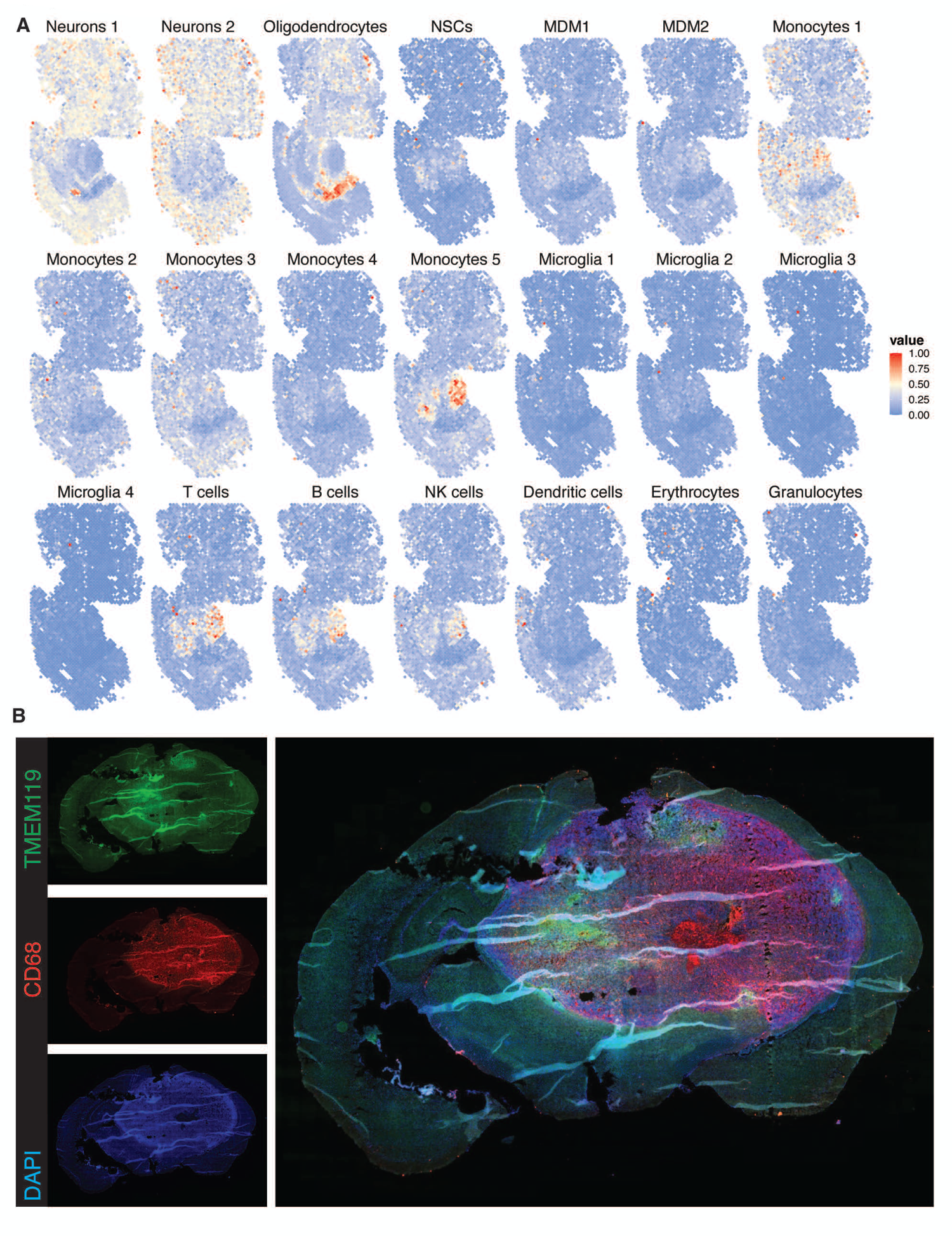
Spatial cellular composition in the brain of mouse GBM. A. Deconvoluted cell type composition in spatial transcriptomics. B. Representative IF image of mouse brain implanted with syngeneic tumor cells stained for TMEM119 and CD68.

**Supplementary Figure 5.**
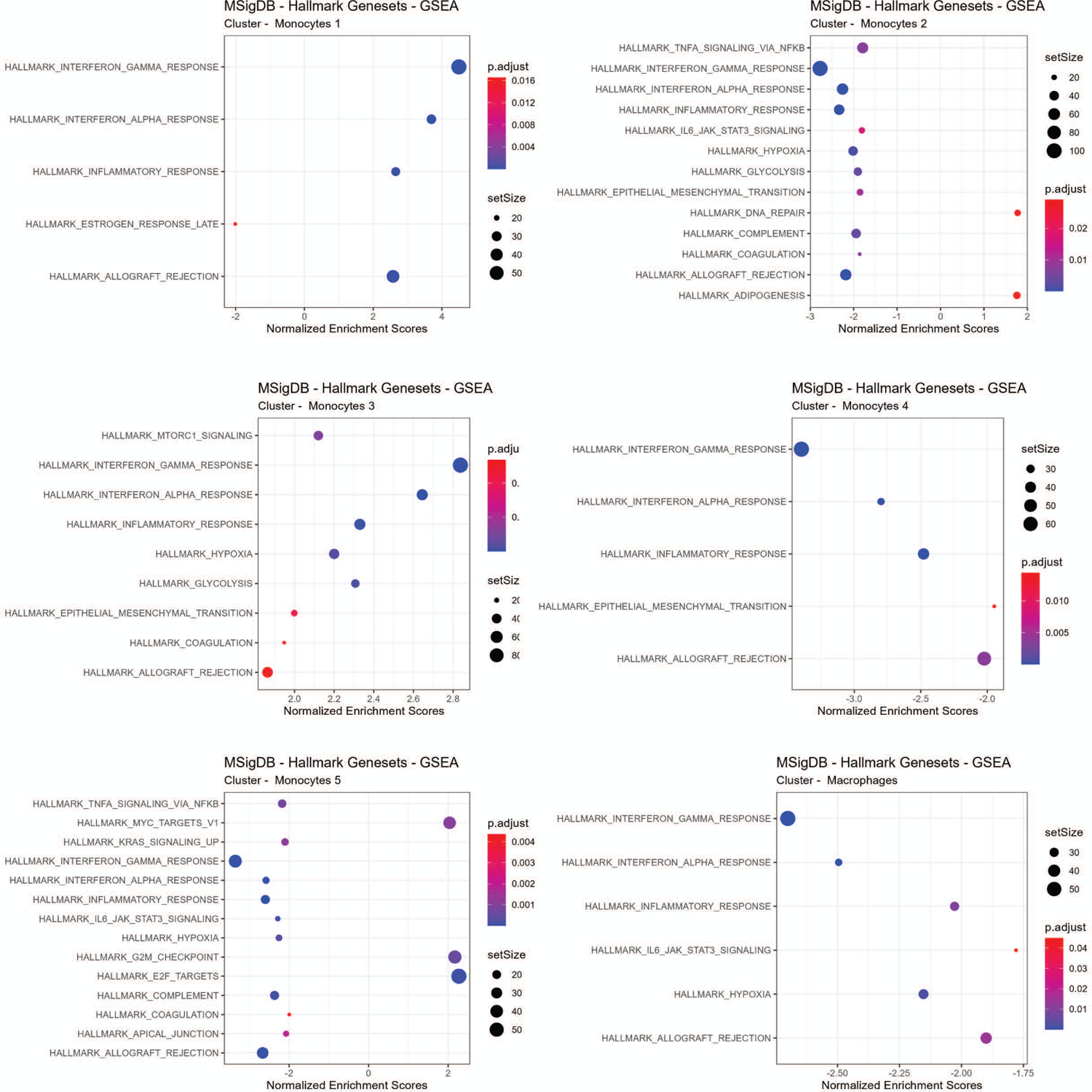
Gene set enrichment analysis of hallmark of cancer gene sets for monocyte subsets from scRNA-seq of mouse GBM.

**Supplementary Figure 6.**
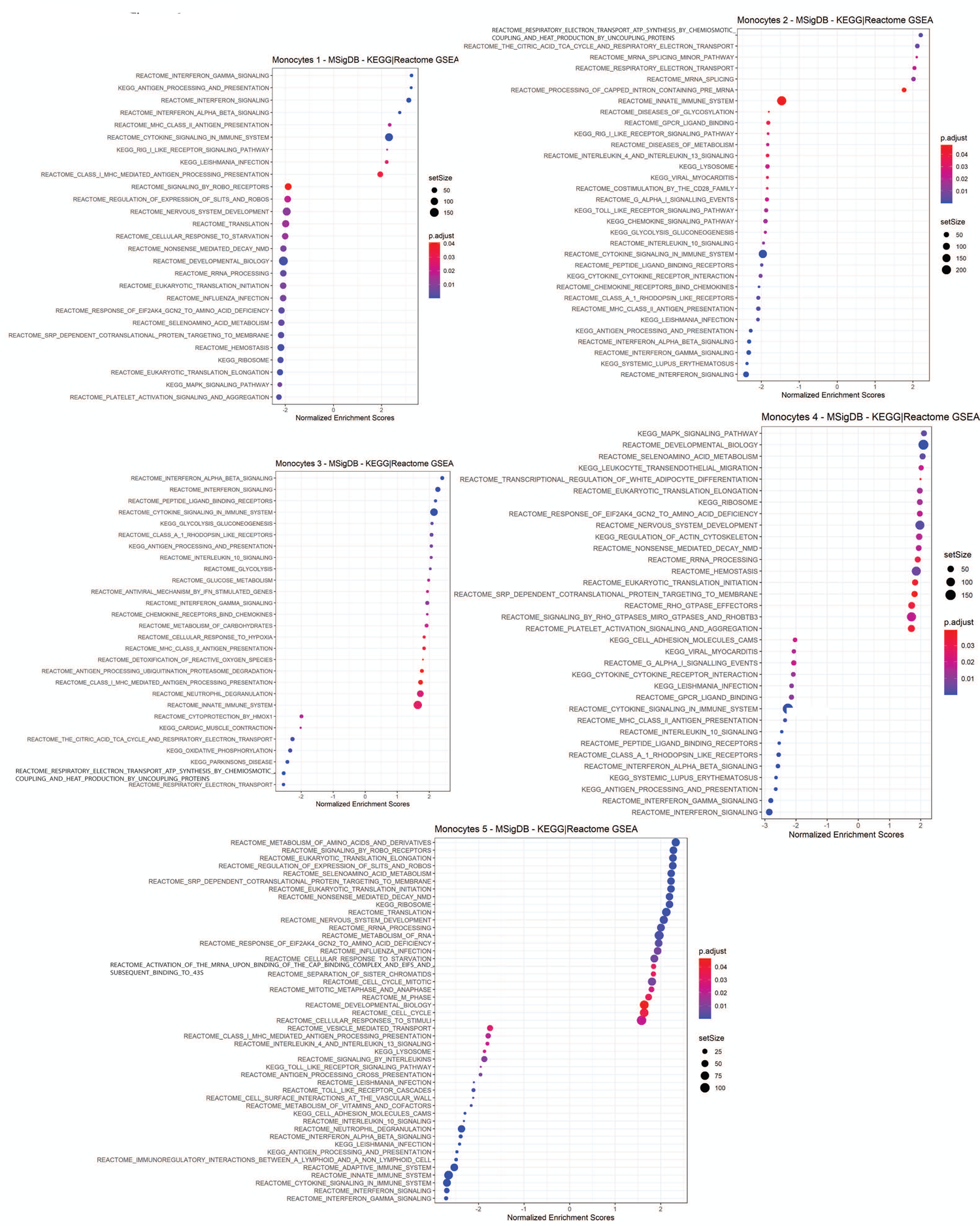
Gene set enrichment analysis of KEGG, and REACTOME for monocyte subsets from scRNA-seq of mouse GBM.

**Supplementary Figure 7.**
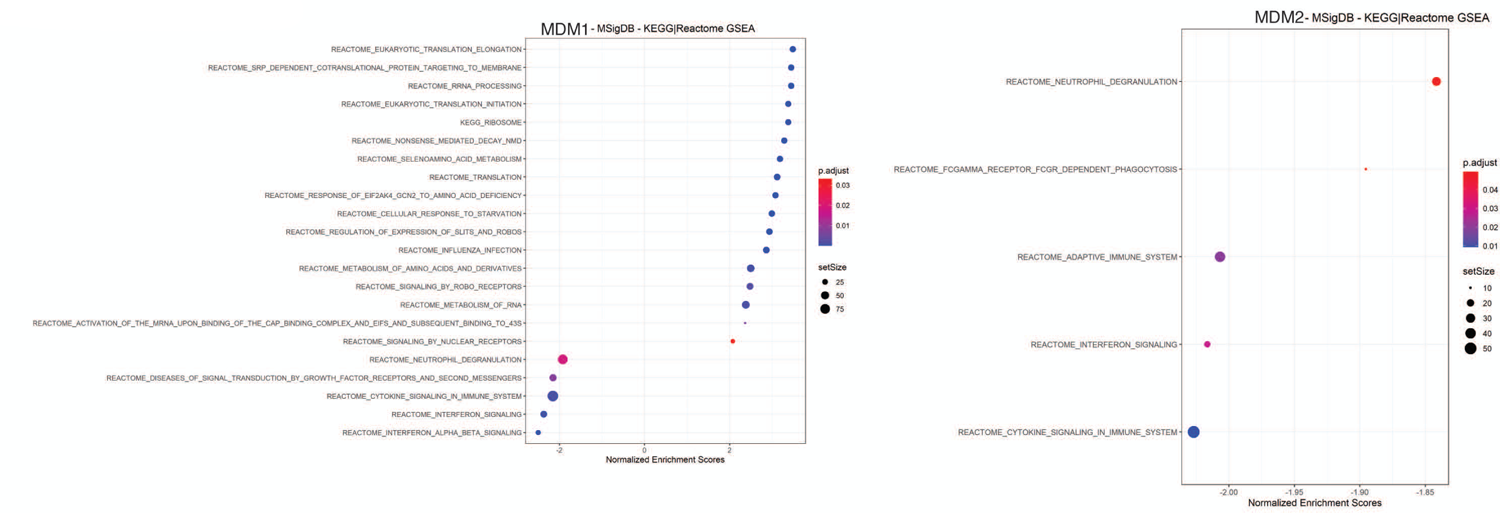
Gene set enrichment analysis of KEGG, and REACTOME for MDM subsets from scRNA-seq of mouse GBM.

**Supplementary Figure 8.**
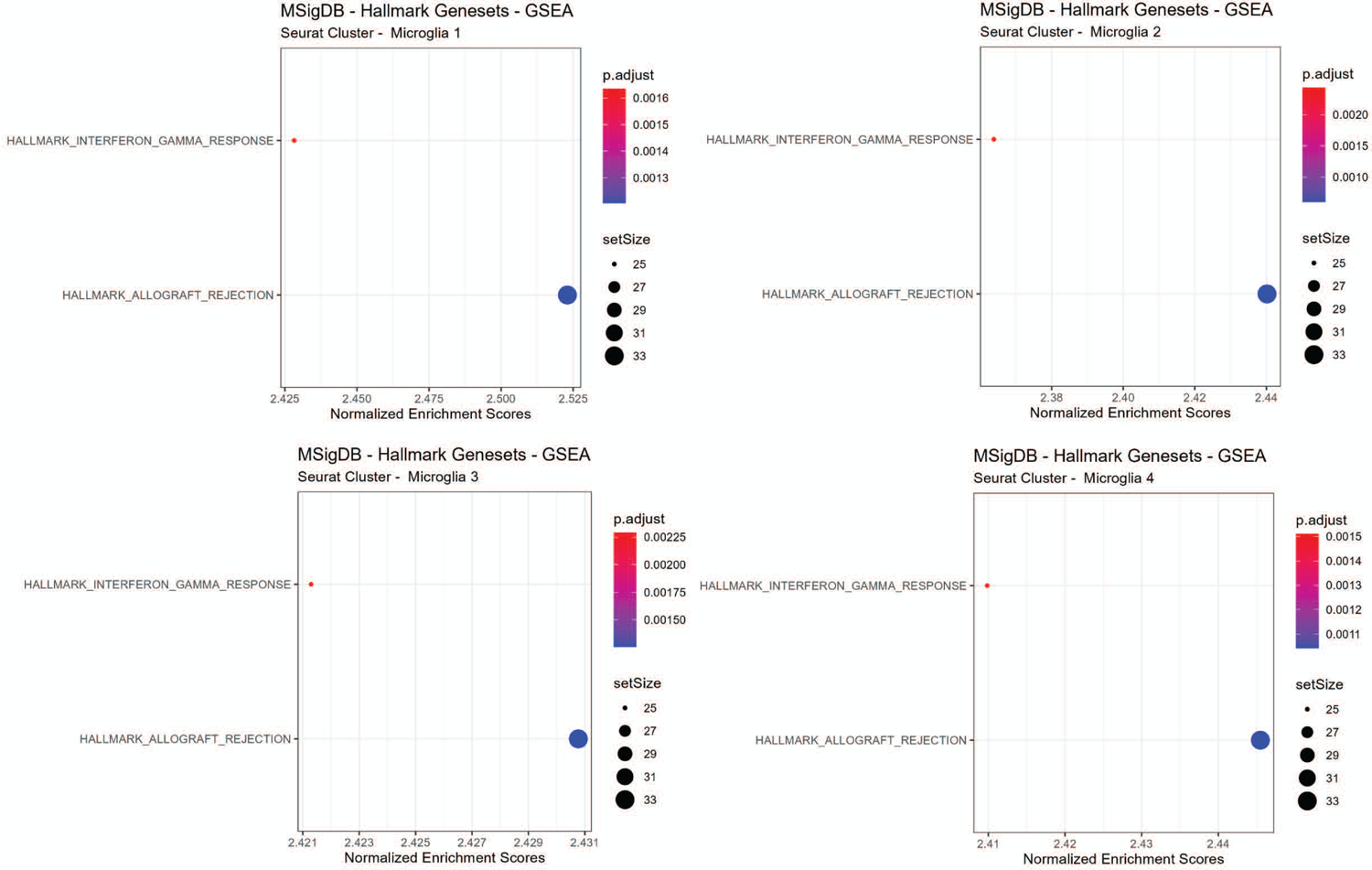
Gene set enrichment analysis of hallmark of cancer gene sets for MG subsets from scRNA-seq of mouse GBM.

**Supplementary Figure 9.**
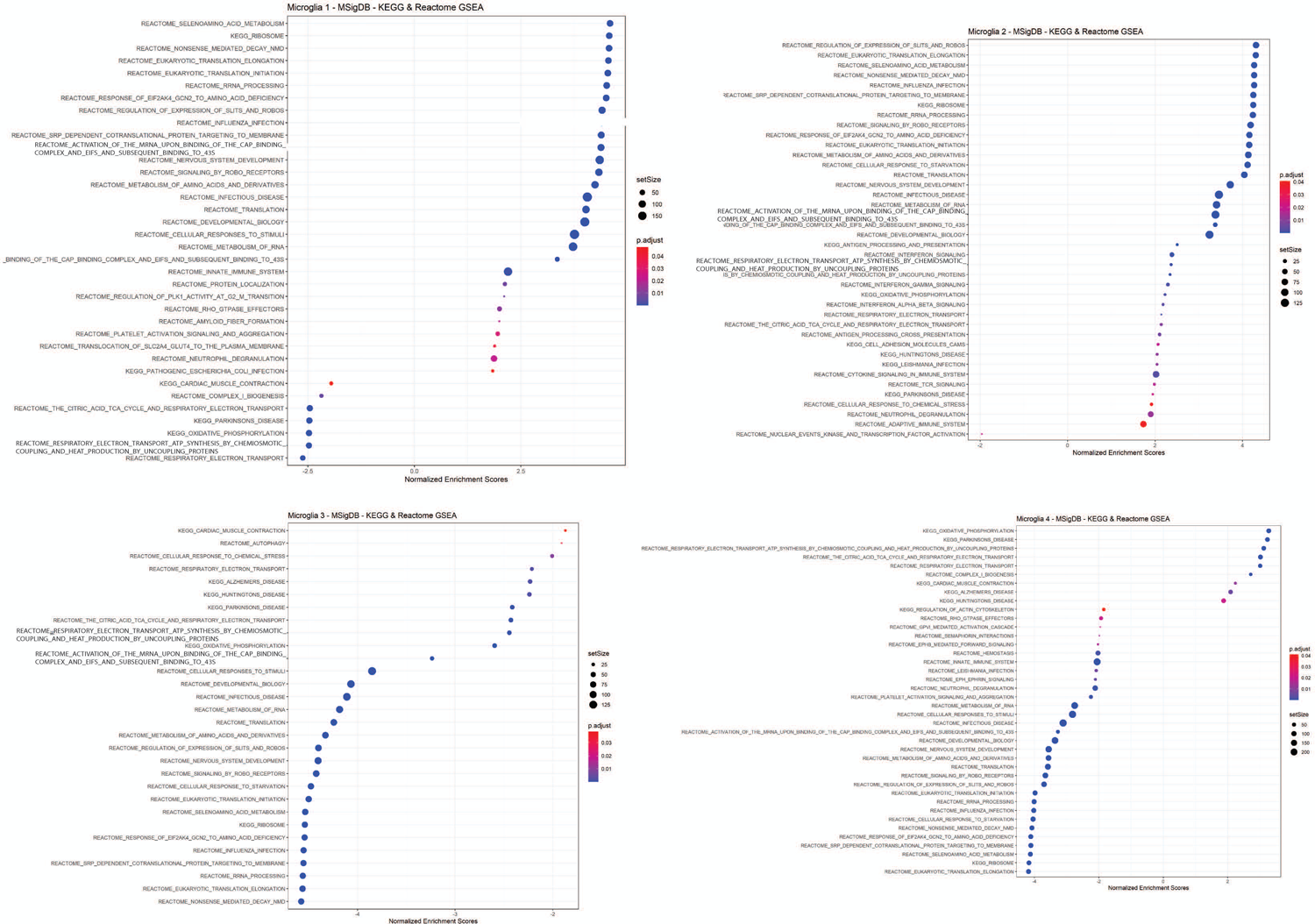
Gene set enrichment analysis of KEGG, and REACTOME for MG subsets from scRNA-seq of mouse GBM.

**Supplementary Figure 10.**
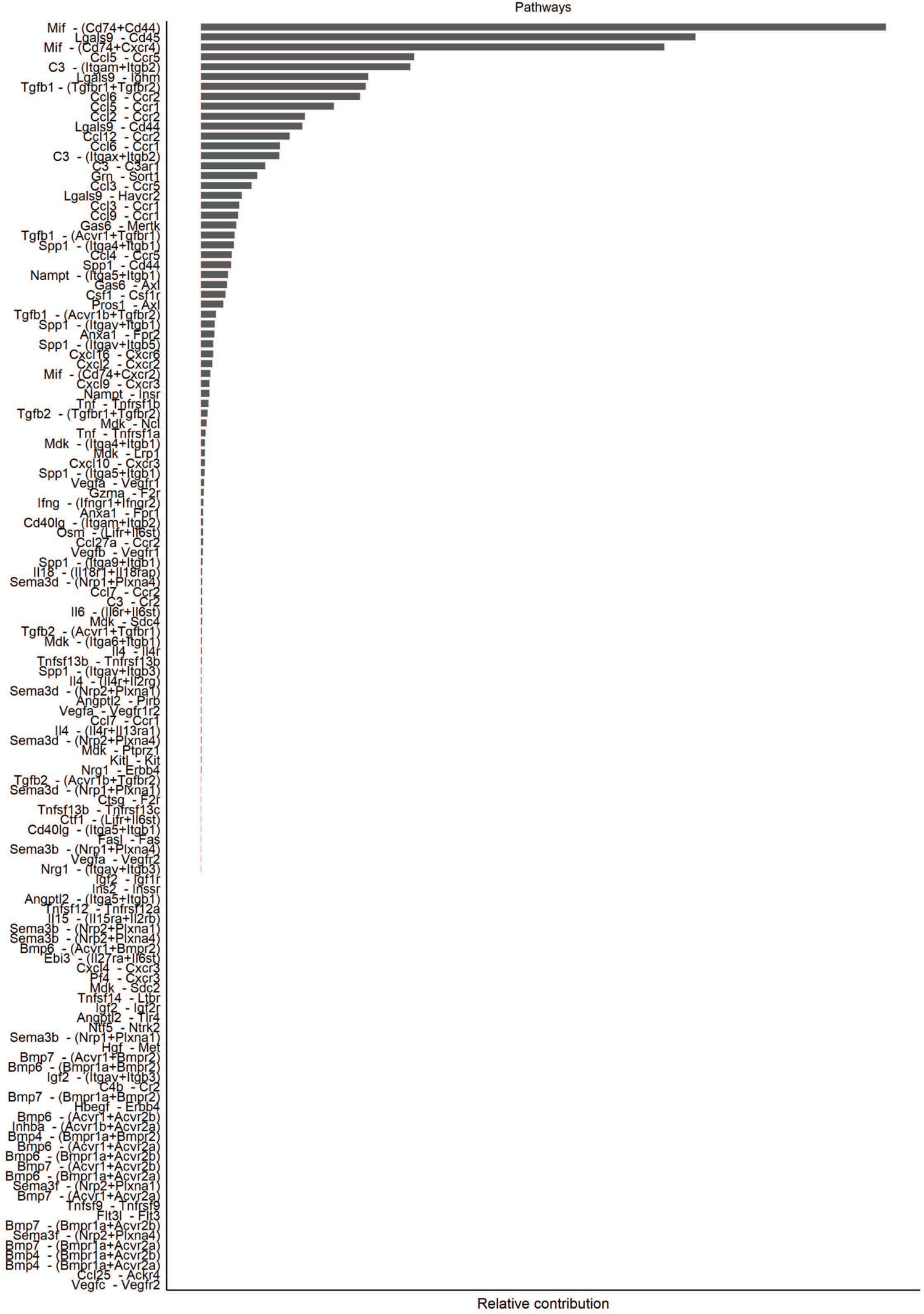
Signaling network in the GBM microenvironment. Contribution of each ligand-receptor pair to the overall signaling pathway calculated in scRNA-seq of mouse GBM.

**Supplementary Figure 11.**
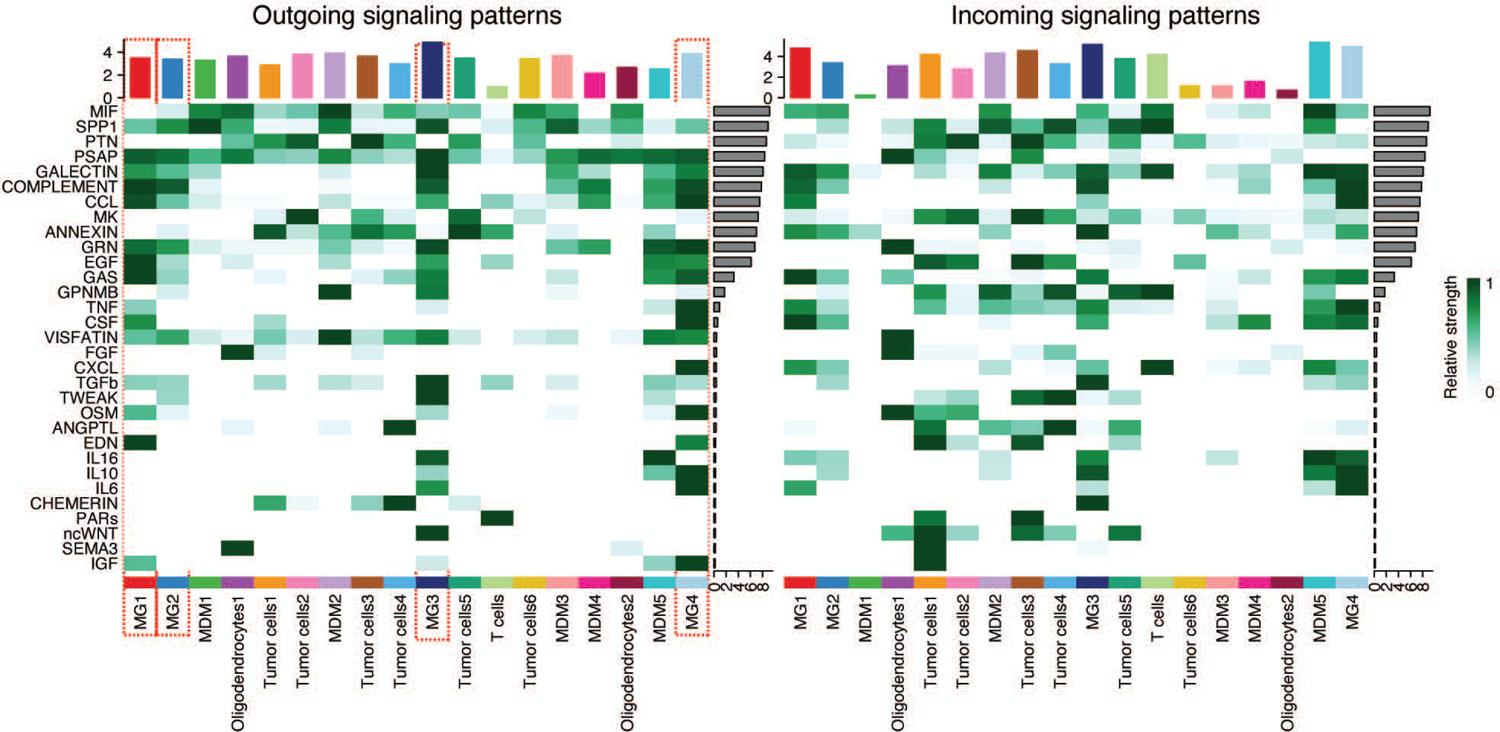
Network contributing to outgoing or incoming signaling in human GBM. Heatmaps showing signals contributing most to outgoing or incoming signaling of certain cell groups, respectively in the scRNA-seq of human GBM.

**Supplementary Figure 12.**
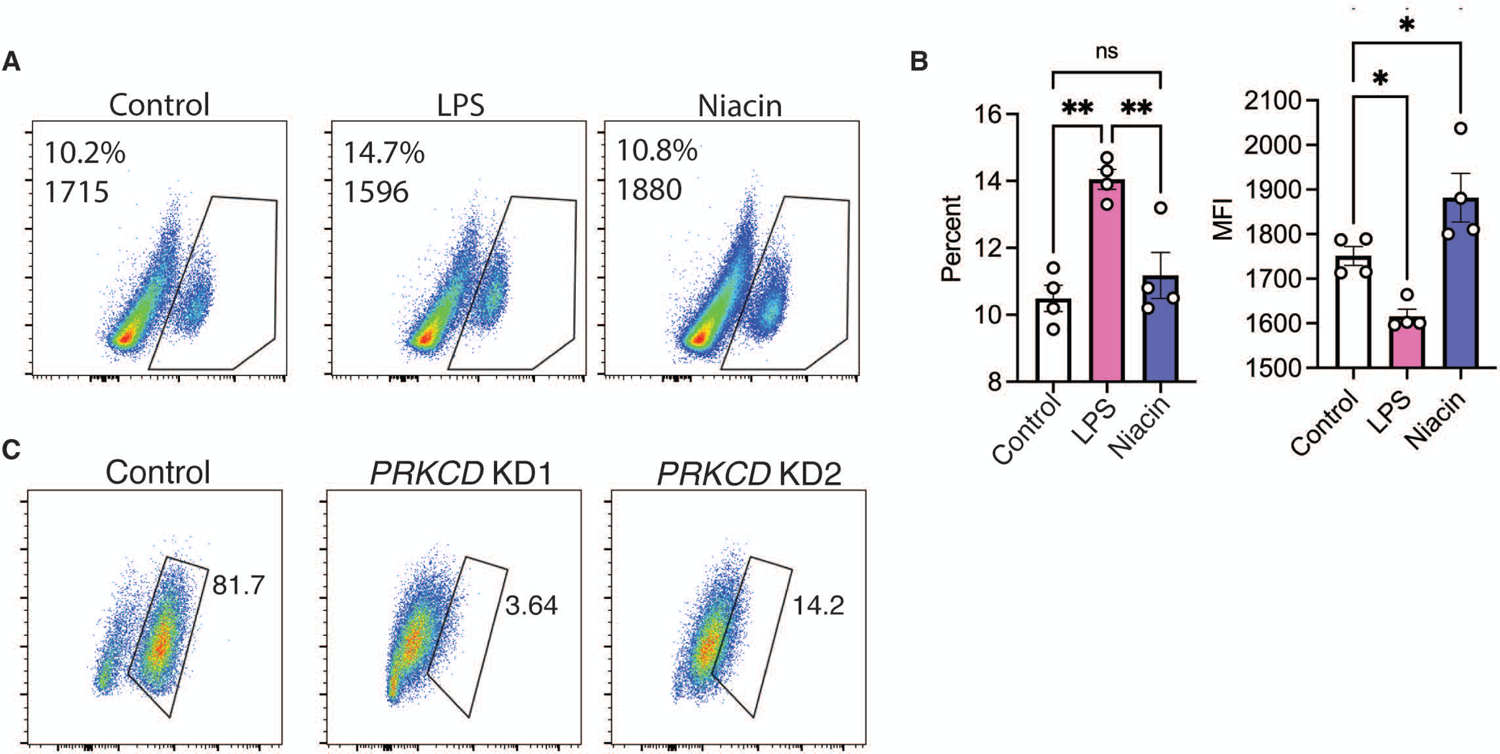
Involvement of PKCδ in the phagocytosis of tumor cells. A. Representative flow cytometry plots of phagocytosis of pHrodo Bioparticles in human primary MG treated with LPS and niacin. B. Quantification of phagocytosis in human MG. C. Flow cytometry plots showing downregulation of *PRKCD* using two different shRNAs in CHME cells.

**Supplementary Figure 13.**
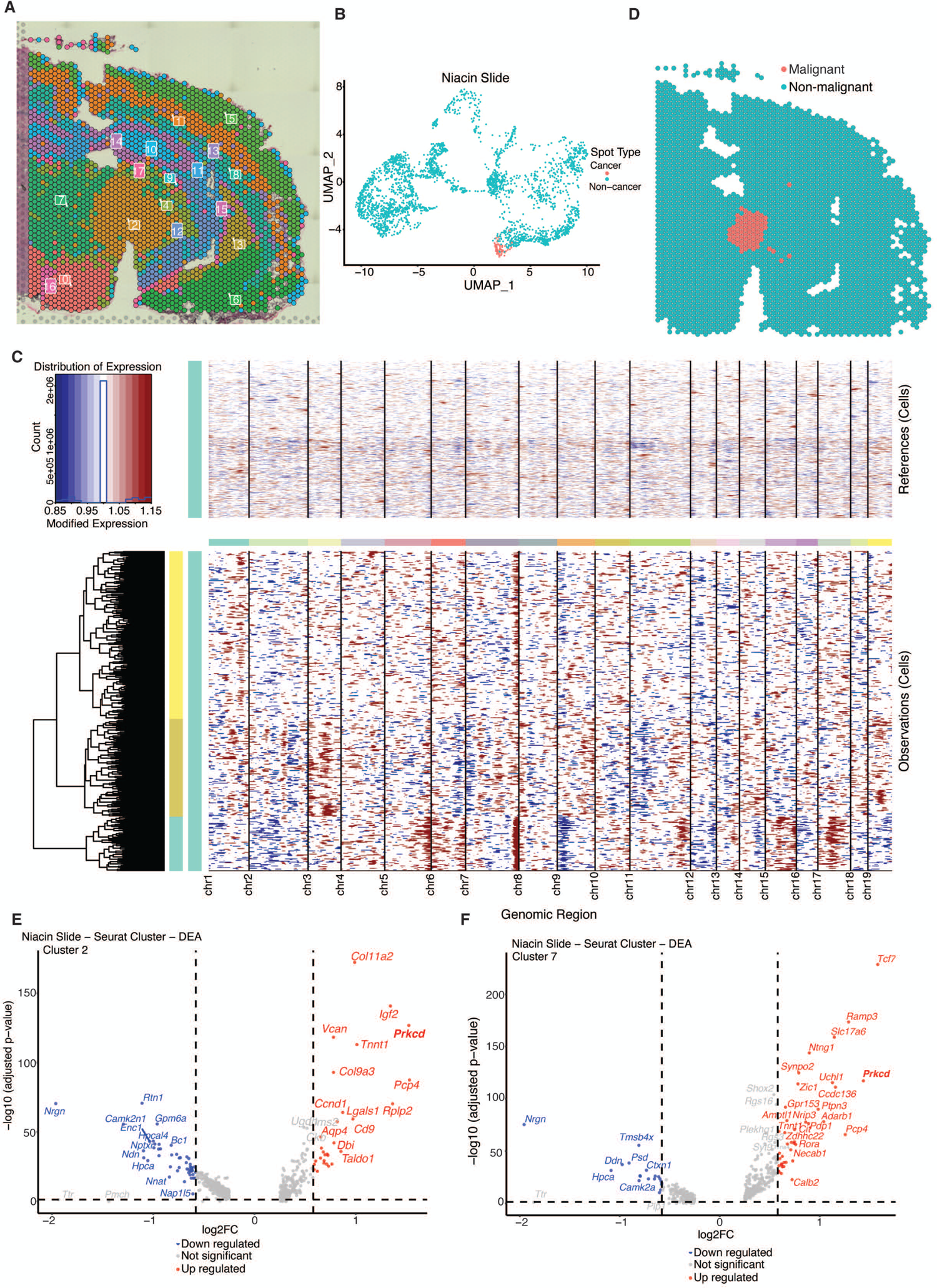
Niacin treated spatial transcriptomic slide. A. Spatial Dimplot of Seurat clusters overlaid on tissue images of tumor mice treated with niacin. B. InferCNV analysis of spatial clusters highlighting malignant and non-malignant spots on tissue images. C. InferCNV heatmap showing CNV events across different chromosomes in cancerous spots (bottom) and normal spots (top). D. Spatial distribution depicting malignant and non-malignant areas. E-F. Volcano plots of DEGs between Seurat clusters for cluster 2 and cluster 7.

**Supplementary Figure 14.**
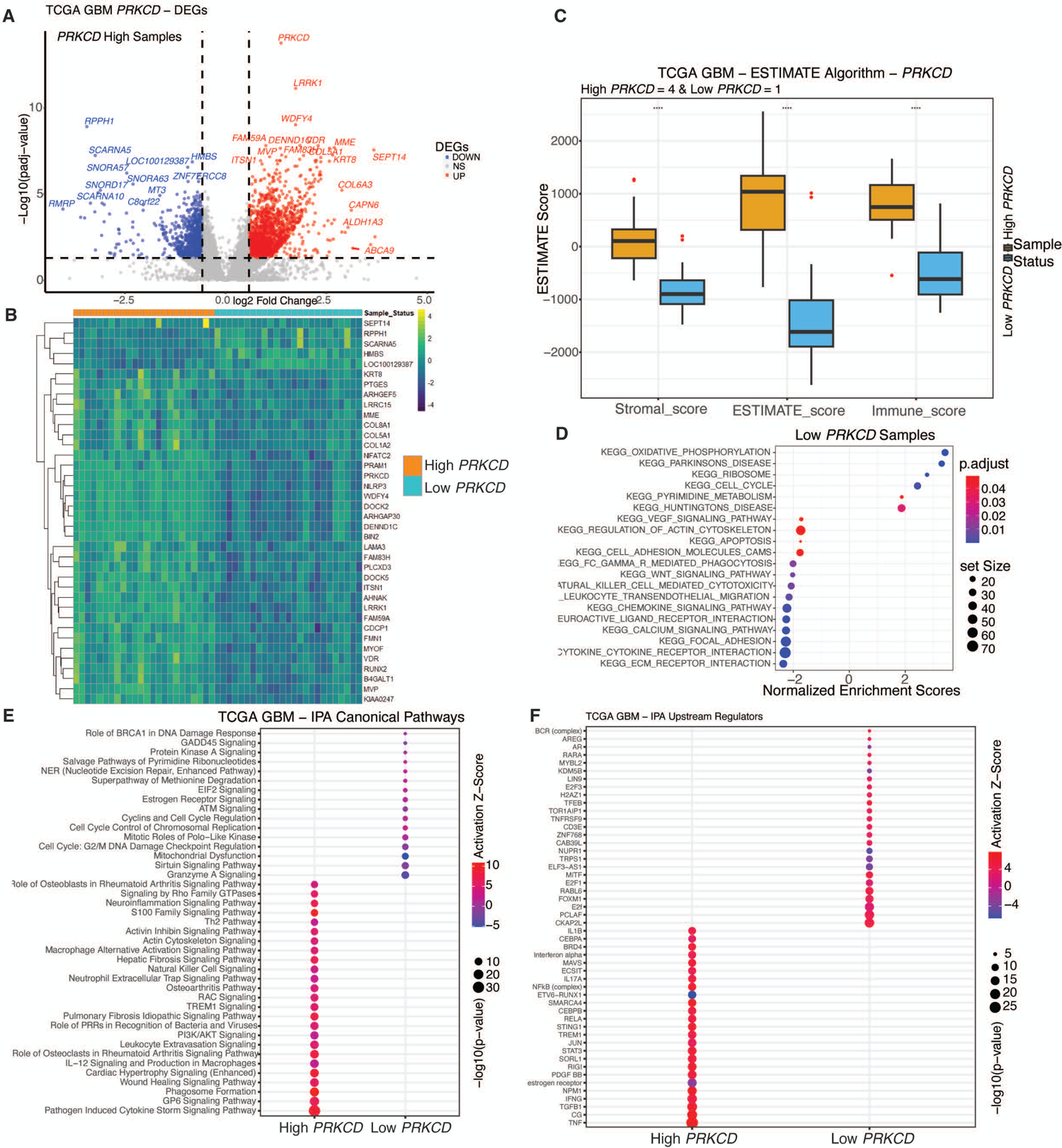
Analysis of TCGA database for *PRKCD* expression. A. Volcano plot of DEGs between *PRKCD* high versus low samples using TCGA GBM RNA samples. B. Heatmap of top DEGs showing difference in gene counts between sample conditions. C. ESTIMATE analysis examining immune, stromal, and purity scores between *PRKCD* high and low samples, significant differences between conditions determined using Wilcoxon rank test (p < 0.05). D. GSEA and MSigDB analysis of KEGG and REACTOME pathways in *PRKCD* low samples. E-F. Dot plots of IPA results on canonical pathways and upstream regulators enriched in either *PRKCD* high or low samples.

## Notes

### Competing Interest Statement

The authors have declared no competing interest.

### Summary of Updates

The new version of the manuscript includes additional supplementary data that supports the main findings of the study. Furthermore, both the manuscript and figures have undergone improvements, providing a clearer narrative that aids readers in better understanding the data and conclusions.

## References

1. Stupp R, Mason WP, van den Bent MJ, Weller M, Fisher B, Taphoorn MJ, et al. Radiotherapy plus concomitant and adjuvant temozolomide for glioblastoma. N Engl J Med 2005;352:987–96

2. Stupp R, Taillibert S, Kanner A, Read W, Steinberg D, Lhermitte B, et al. Effect of Tumor-Treating Fields Plus Maintenance Temozolomide vs Maintenance Temozolomide Alone on Survival in Patients With Glioblastoma: A Randomized Clinical Trial. JAMA 2017;318:2306–16

3. Klemm F, Maas RR, Bowman RL, Kornete M, Soukup K, Nassiri S, et al. Interrogation of the Microenvironmental Landscape in Brain Tumors Reveals Disease-Specific Alterations of Immune Cells. Cell 2020;181:1643–60.e17

4. Richards LM, Whitley OKN, MacLeod G, Cavalli FMG, Coutinho FJ, Jaramillo JE, et al. Gradient of Developmental and Injury Response transcriptional states defines functional vulnerabilities underpinning glioblastoma heterogeneity. Nat Cancer 2021;2:157–73

5. Mirzaei R, D’Mello C, Liu M, Nikolic A, Kumar M, Visser F, et al. Single-Cell Spatial Analysis Identifies Regulators of Brain Tumor-Initiating Cells. Cancer Res 2023;83:1725–41

6. LeBlanc VG, Trinh DL, Aslanpour S, Hughes M, Livingstone D, Jin D, et al. Single-cell landscapes of primary glioblastomas and matched explants and cell lines show variable retention of inter- and intratumor heterogeneity. Cancer Cell 2022

7. Mirzaei R, Yong VW. Microglia-T cell conversations in brain cancer progression. Trends Mol Med 2022;28:951–63

8. Yeo AT, Rawal S, Delcuze B, Christofides A, Atayde A, Strauss L, et al. Single-cell RNA sequencing reveals evolution of immune landscape during glioblastoma progression. Nat Immunol 2022;23:971–84

9. González-Silva L, Quevedo L, Varela I. Tumor Functional Heterogeneity Unraveled by scRNA-seq Technologies. Trends Cancer 2020;6:13–9

10. Wischnewski V, Maas RR, Aruffo PG, Soukup K, Galletti G, Kornete M, et al. Phenotypic diversity of T cells in human primary and metastatic brain tumors revealed by multiomic interrogation. Nat Cancer 2023;4:908–24

11. Armingol E, Officer A, Harismendy O, Lewis NE. Deciphering cell-cell interactions and communication from gene expression. Nat Rev Genet 2021;22:71–88

12. Laviron M, Petit M, Weber-Delacroix E, Combes AJ, Arkal AR, Barthélémy S, et al. Tumor-associated macrophage heterogeneity is driven by tissue territories in breast cancer. Cell Rep 2022;39:110865

13. Ravi VM, Will P, Kueckelhaus J, Sun N, Joseph K, Salié H, et al. Spatially resolved multi-omics deciphers bidirectional tumor-host interdependence in glioblastoma. Cancer Cell 2022;40:639–55.e13

14. Moldoveanu D, Ramsay L, Lajoie M, Anderson-Trocme L, Lingrand M, Berry D, et al. Spatially mapping the immune landscape of melanoma using imaging mass cytometry. Sci Immunol 2022;7:eabi5072

15. Reilly KM, Loisel DA, Bronson RT, McLaughlin ME, Jacks T. Nf1;Trp53 mutant mice develop glioblastoma with evidence of strain-specific effects. Nat Genet 2000;26:109–13

16. Pisklakova A, McKenzie B, Zemp F, Lun X, Kenchappa RS, Etame AB, et al. M011L-deficient oncolytic myxoma virus induces apoptosis in brain tumor-initiating cells and enhances survival in a novel immunocompetent mouse model of glioblastoma. Neuro Oncol 2016;18:1088–98

17. Ren Y, Huang Z, Zhou L, Xiao P, Song J, He P, et al. Spatial transcriptomics reveals niche-specific enrichment and vulnerabilities of radial glial stem-like cells in malignant gliomas. Nat Commun 2023;14:1028

18. Li T, Fan J, Wang B, Traugh N, Chen Q, Liu JS, et al. TIMER: A Web Server for Comprehensive Analysis of Tumor-Infiltrating Immune Cells. Cancer Res 2017;77:e108–e10

19. Li T, Fu J, Zeng Z, Cohen D, Li J, Chen Q, et al. TIMER2.0 for analysis of tumor-infiltrating immune cells. Nucleic Acids Res 2020;48:W509–W14

20. Rybicka JM, Balce DR, Khan MF, Krohn RM, Yates RM. NADPH oxidase activity controls phagosomal proteolysis in macrophages through modulation of the lumenal redox environment of phagosomes. Proc Natl Acad Sci U S A 2010;107:10496–501

21. Sarkar S, Yang R, Mirzaei R, Rawji K, Poon C, Mishra MK, et al. Control of brain tumor growth by reactivating myeloid cells with niacin. Sci Transl Med 2020;12

22. Rawji KS, Young AMH, Ghosh T, Michaels NJ, Mirzaei R, Kappen J, et al. Niacin-mediated rejuvenation of macrophage/microglia enhances remyelination of the aging central nervous system. Acta Neuropathol 2020;139:893–909

23. Harikumar KB, Kunnumakkara AB, Ochi N, Tong Z, Deorukhkar A, Sung B, et al. A novel small-molecule inhibitor of protein kinase D blocks pancreatic cancer growth in vitro and in vivo. Mol Cancer Ther 2010;9:1136–46

24. Lin R, Zhou Y, Yan T, Wang R, Li H, Wu Z, et al. Directed evolution of adeno-associated virus for efficient gene delivery to microglia. Nat Methods 2022;19:976–85

25. Soh JW, Lee EH, Prywes R, Weinstein IB. Novel roles of specific isoforms of protein kinase C in activation of the c-fos serum response element. Mol Cell Biol 1999;19:1313–24

26. Jain S, Rick JW, Joshi RS, Beniwal A, Spatz J, Gill S, et al. Single-cell RNA sequencing and spatial transcriptomics reveal cancer-associated fibroblasts in glioblastoma with protumoral effects. J Clin Invest 2023;133

27. Guilliams M, Bonnardel J, Haest B, Vanderborght B, Wagner C, Remmerie A, et al. Spatial proteogenomics reveals distinct and evolutionarily conserved hepatic macrophage niches. Cell 2022;185:379–96.e38

28. Sorin M, Rezanejad M, Karimi E, Fiset B, Desharnais L, Perus LJM, et al. Single-cell spatial landscapes of the lung tumour immune microenvironment. Nature 2023;614:548–54

29. Karimi E, Yu MW, Maritan SM, Perus LJM, Rezanejad M, Sorin M, et al. Single-cell spatial immune landscapes of primary and metastatic brain tumours. Nature 2023;614:555–63

30. Phillips D, Matusiak M, Gutierrez BR, Bhate SS, Barlow GL, Jiang S, et al. Immune cell topography predicts response to PD-1 blockade in cutaneous T cell lymphoma. Nat Commun 2021;12:6726

31. Evans KT, Blake K, Longworth A, Coburn MA, Insua-Rodríguez J, McMullen TP, et al. Microglia promote anti-tumour immunity and suppress breast cancer brain metastasis. Nat Cell Biol 2023

32. Kikkawa U, Matsuzaki H, Yamamoto T. Protein kinase C delta (PKC delta): activation mechanisms and functions. J Biochem 2002;132:831–9

33. Parihar SP, Ozturk M, Marakalala MJ, Loots DT, Hurdayal R, Maasdorp DB, et al. Protein kinase C-delta (PKCδ), a marker of inflammation and tuberculosis disease progression in humans, is important for optimal macrophage killing effector functions and survival in mice. Mucosal Immunol 2018;11:496–511

34. Neehus AL, Moriya K, Nieto-Patlán A, Le Voyer T, Lévy R, Özen A, et al. Impaired respiratory burst contributes to infections in PKCδ-deficient patients. J Exp Med 2021;218

35. Migliozzi S, Oh YT, Hasanain M, Garofano L, D’Angelo F, Najac RD, et al. Integrative multi-omics networks identify PKCδ and DNA-PK as master kinases of glioblastoma subtypes and guide targeted cancer therapy. Nat Cancer 2023;4:181–202

36. Sarkar S, Yong VW. Reduction of protein kinase C delta attenuates tenascin-C stimulated glioma invasion in three-dimensional matrix. Carcinogenesis 2010;31:311–7

37. Bessa C, Soares J, Raimundo L, Loureiro JB, Gomes C, Reis F, et al. Discovery of a small-molecule protein kinase Cδ-selective activator with promising application in colon cancer therapy. Cell Death Dis 2018;9:23

38. Liu H, Sun Y, Zhang Q, Jin W, Gordon RE, Zhang Y, et al. Pro-inflammatory and proliferative microglia drive progression of glioblastoma. Cell Rep 2021;36:109718

39. Khan F, Pang L, Dunterman M, Lesniak MS, Heimberger AB, Chen P. Macrophages and microglia in glioblastoma: heterogeneity, plasticity, and therapy. J Clin Invest 2023;133

40. Giuliani F, Hader W, Yong VW. Minocycline attenuates T cell and microglia activity to impair cytokine production in T cell-microglia interaction. J Leukoc Biol 2005;78:135–43

41. Mishra MK, Wang J, Keough MB, Fan Y, Silva C, Sloka S, et al. Laquinimod reduces neuroaxonal injury through inhibiting microglial activation. Ann Clin Transl Neurol 2014;1:409–22

42. Mirzaei R, Gordon A, Zemp FJ, Kumar M, Sarkar S, Luchman HA, et al. PD-1 independent of PD-L1 ligation promotes glioblastoma growth through the NFκB pathway. Sci Adv 2021;7:eabh2148

43. Dzikowski L, Mirzaei R, Sarkar S, Kumar M, Bose P, Bellail A, et al. Fibrinogen in the glioblastoma microenvironment contributes to the invasiveness of brain tumor-initiating cells. Brain Pathol 2021;31:e12947

44. Mirzaei R, Sarkar S, Dzikowski L, Rawji KS, Khan L, Faissner A, et al. Brain tumor-initiating cells export tenascin-C associated with exosomes to suppress T cell activity. Oncoimmunology 2018;7:e1478647

45. Kelly JJ, Stechishin O, Chojnacki A, Lun X, Sun B, Senger DL, et al. Proliferation of human glioblastoma stem cells occurs independently of exogenous mitogens. Stem Cells 2009;27:1722–33

46. Butler A, Hoffman P, Smibert P, Papalexi E, Satija R. Integrating single-cell transcriptomic data across different conditions, technologies, and species. Nat Biotechnol 2018;36:411–20

47. Jiang W, Hua R, Wei M, Li C, Qiu Z, Yang X, et al. An optimized method for high-titer lentivirus preparations without ultracentrifugation. Sci Rep 2015;5:13875

48. Grace PM, Strand KA, Galer EL, Urban DJ, Wang X, Baratta MV, et al. Morphine paradoxically prolongs neuropathic pain in rats by amplifying spinal NLRP3 inflammasome activation. Proc Natl Acad Sci U S A 2016;113:E3441–50

49. Challis RC, Ravindra Kumar S, Chan KY, Challis C, Beadle K, Jang MJ, et al. Systemic AAV vectors for widespread and targeted gene delivery in rodents. Nat Protoc 2019;14:379–414

50. Danaher P, Warren S, Dennis L, D’Amico L, White A, Disis ML, et al. Gene expression markers of Tumor Infiltrating Leukocytes. J Immunother Cancer 2017;5:18

51. Ochocka N, Segit P, Walentynowicz KA, Wojnicki K, Cyranowski S, Swatler J, et al. Single-cell RNA sequencing reveals functional heterogeneity of glioma-associated brain macrophages. Nat Commun 2021;12:1151

52. Rajendran S, Hu Y, Canella A, Peterson C, Gross A, Cam M, et al. Single-cell RNA sequencing reveals immunosuppressive myeloid cell diversity during malignant progression in a murine model of glioma. Cell Rep 2023;42:112197

